# Ifnar1 signaling breaks the hepatic urea cycle to regulate adaptive immunity

**DOI:** 10.1101/762310

**Authors:** Alexander Lercher, Anannya Bhattacharya, Alexandra M. Popa, Michael Caldera, Moritz F. Schlapansky, Hatoon Baazim, Peter Majek, Julia S. Brunner, Lindsay J. Kosack, Dijana Vitko, Theresa Pinter, Bettina Gürtl, Daniela Reil, Ulrich Kalinke, Keiryn L. Bennett, Jörg Menche, Paul N. Cheng, Gernot Schabbauer, Michael Trauner, Kristaps Klavins, Andreas Bergthaler

## Abstract

Infections induce complex host responses linked to antiviral defense, inflammation and tissue damage and repair. These processes are increasingly understood to involve systemic metabolic reprogramming. We hypothesized that the liver as a central metabolic hub may orchestrate many of these changes during infection. Thus, we investigated the systemic interplay between inflammation and metabolism in a mouse model of chronic viral infection and hepatitis. Here we show that virus-induced type I interferon (IFN-I) modulates wide-spread metabolic alterations of the liver in a hepatocyte-intrinsic Ifnar1-dependent way. Specifically, IFN-I repressed the transcription of numerous genes with metabolic function including *Otc* and *Ass1*, which encode enzymes of the urea cycle. This led to decreased arginine and increased ornithine concentrations in the circulation, resulting in suppressed virus-specific CD8 T cell responses and ameliorated liver pathology. These findings establish IFN-I-induced modulation of hepatic metabolism and the urea cycle as an endogenous mechanism of immunoregulation.

## Introduction

Pathogens and concurrent tissue damage elicit complex context-dependent inflammatory response programs (Hotamisligil, 2017). Chronic infections represent a particular challenge for the host organism, which is exposed to prolonged inflammation that may predispose to various co-morbidities such as susceptibility to secondary infections and cancer (Okin and Medzhitov, 2012; Rehermann and Nascimbeni, 2005; Virgin et al., 2009). The same inflammatory pathways can also control cellular tissue homeostasis and metabolism (Kotas and Medzhitov, 2015; Medzhitov, 2008; O’Neill and Pearce, 2016). Different cell types and organs communicate with each other through soluble cytokines, thereby determining the quality, magnitude and duration of both local and systemic immune responses (Medzhitov, 2008). The liver is a central metabolic organ but also represents an immunoregulatory hub between blood-borne pathogens and the immune system (Jenne and Kubes, 2013; Protzer et al., 2012; Racanelli and Rehermann, 2006). Hepatocytes are the functional unit of the liver parenchyma and the most abundant cell type of the liver. As such their principal task is the turnover of metabolites during homeostasis. Yet they are also important immune signaling platforms that produce and react to a range of cytokines upon inflammation (Crispe, 2016; Racanelli and Rehermann, 2006; Zhou et al., 2015). Hepatocytes themselves are permissible to a variety of chronic viruses including hepatitis B virus (HBV) and hepatitis C virus (HCV) in humans and lymphocytic choriomeningitis virus (LCMV) in mice (Guidotti et al., 1999; Rehermann and Nascimbeni, 2005). The chronic infection model of LCMV represents a well-established and pathophysiologically relevant experimental model for the study of host-pathogen interactions and immune responses that induces a vigorous CD8 T cell-dependent hepatitis (Zehn and Wherry, 2015; Zinkernagel et al., 1986). The involved immunopathologic mechanisms seen in the LCMV model are similar to those observed in patients chronically infected with HBV or HCV (Guidotti et al., 1999; Rehermann and Nascimbeni, 2005).

Soluble inflammatory signals act mainly through cytokine receptors (Hotamisligil, 2017; Protzer et al., 2012; Racanelli and Rehermann, 2006). Type I interferons (IFN-I) are central antiviral cytokines that signal through the ubiquitously expressed IFNAR receptor, which is comprised of the two subunits IFNAR1 and IFNAR2. This induces the expression of a broad array of genes described as interferon-stimulated genes (ISGs). ISGs exert antiviral functions by direct interference with viral replication and immunoregulatory properties (McNab et al., 2015; Schoggins et al., 2011). More recently, IFN-I are also recognized as modulators of metabolism, such as cellular lipid metabolism and redox homeostasis (Bhattacharya et al., 2015; Pantel et al., 2014; Wu et al., 2016; York et al., 2015). Cytokine-induced regulation of liver metabolism is expected to result in altered metabolite turnover and release that impacts distal organs (van den Berghe, 1991; Norata et al., 2015). Immune cells and in particular T cells critically depend on certain metabolites to efficiently perform their functions and are, thus, susceptible to altered metabolite availability (Chang et al., 2015; Geiger et al., 2016; Johnson et al., 2018; Pearce et al., 2009). In line with this, a frequent immune evasion mechanism of cancer is the depletion of essential amino acids or glucose in the tumor microenvironment (Buck et al., 2017; Chang et al., 2015; Ma et al., 2017; Murray, 2015).

In this study we investigated the chronic infection model of LCMV together using an unbiased integrative approach to unveil inflammation-driven endogenous regulation of liver metabolism and study its impact on the systemic immune response and tissue pathology.

## Results

### Identification of inflammatory-metabolic changes in the liver during chronic infection

We infected C57BL/6J wildtype (WT) mice with 2×10^6^ FFU of the chronic strain clone 13 of LCMV and quantified infectious virus particles and viral RNA in the liver (Figure 1A). This confirmed the peak of viral propagation around day 8 after infection with a subsequent decline of viral loads. Liver damage was assessed by the clinical hallmark parameters alanine aminotransferase (ALT) and aspartate aminotransferase (AST) and peaked on day 8 to 12 after infection (Figure 1B). In an unbiased approach to virus-induced changes in the liver, we collected liver tissue from infected mice at different phases of infection (day 2 ≈ innate phase, day 8 ≈ peak of disease, day 30 ≈ chronic phase, day 60 ≈ resolving phase) and performed transcriptome analyses by RNA-seq. The transcriptomic data showed high intra-replica reproducibility and individual samples clustered according to the time course of infection (Figure 1C), highlighting phase-specific changes in gene expression and the gradual recovery of mice by 60 days after infection. In total, 3626 transcripts were differentially regulated (Table S1A) and the most differentially expressed genes were found on day 2 and 8 after infection (Figure S1A). Hierarchical clustering identified three broad categories of gene expression programs, including transcripts that were found already regulated on day 2 (cluster 1 to 4) or induced on day 8 (cluster 5 to 10) as well as transcripts that were repressed during infection (cluster 11 to 15) (Figure 1D and Table S2A). To corroborate transcriptional alterations, we performed tissue proteomics and quantified 5586 proteins in the liver (Table S1B). Protein changes correlated with differentially expressed transcripts (Figure 1E, S1A and Table S2C). Of the identified clusters, differentially expressed genes associated with antiviral IFN-I signaling were found to be enriched on day 2 (clusters 1 to 4, Figure 1F and Table S2B) while leukocyte-associated genes were mainly found on day 8 after infection (clusters 5 to 10, Figure 1F and Table S2B). Some of these differential expression changes are likely to reflect hepatic immune cell infiltration (Figure S1D).

**Figure 1:**
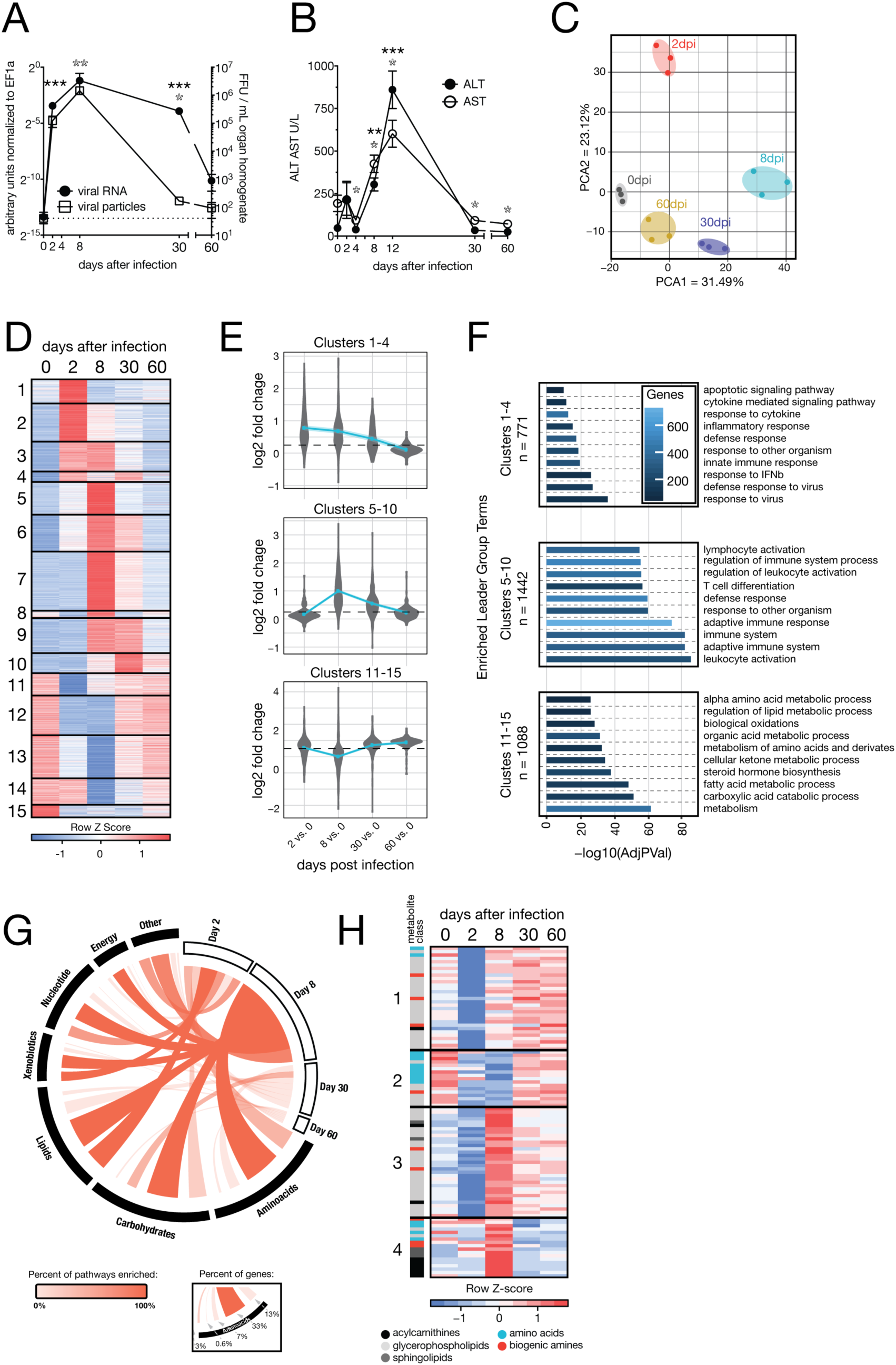
LCMV Cl13 induces hepatic metabolic reprogramming translating to changes in systemic metabolism during the course of infection. (A) Viremia and RNemia of LCMV clone 13 infected liver tissue. (B) Serum alanine transferase (ALT) and aspartate aminotransferase (AST) levels upon LCMV clone 13 infection. (C) Principal component analyses (PCA) of transcriptomic changes in liver tissue. (D) Hierarchical clustering (FPKM, k-means, Pearson’s correlation) of significantly changed genes. (E) Regulation of corresponding detected proteins. (F) Enriched GO terms and pathways (ClueGO) in the clusters identified in (D). (G) Regulated metabolic transcripts and proteins enriched on KEGG database. (H) Hierarchical clustering of significantly regulated serum metabolites (k-means, Pearson’s correlation). Symbols represent the arithmetic mean ±S.E.M. ns = not significant * P < 0.05 ** P < 0.01 *** P < 0.001 (Student’s t-test).

Intriguingly, our enrichment analyses indicated wide-spread metabolic reprogramming of liver tissue during LCMV infection (Figure S1D) with particularly pronounced effects seen for downregulated metabolic pathways relating to lipid- and amino acid metabolism on day 2 and 8 after infection (clusters 11 to 15, Figure 1F and Table S2B). For an in-depth integration of these metabolic changes, we performed enrichment analyses of the union of differentially regulated transcripts and proteins at each stage of viral infection on KEGG metabolic pathways. We identified global reprogramming of processes related to all the major classes of metabolites.

Modulation of hepatic pathways involving lipids and essential amino acids was initiated on day 2 and peaking on day 8 after infection (Figure 1G and Table S1C). Differentially expressed metabolic processes were also found in the chronic and resolving phases of infection, although the number of modulated transcripts and proteins were lower (Figures 1G, S1A and S1B).

To address the effects of virus-induced alterations on the metabolic output of the liver, we performed targeted metabolomics of serum, focusing on amino acids, biogenic amines, sphingolipids, acylcarnitines and glycerophospholipids. We found 99 of 180 metabolites to be significantly regulated at one or more time points (Figure 1H and Table S3A). Downregulation of metabolites on day 2 (Figure S1E) was observed for glycerophospholipids and sphingolipids (Figure 1H, clusters 1 and 3) while acylcarnitines (Figure 1H, cluster 4) were rather upregulated on day 8 after infection (Figure 1H). Systemic amino acids and biogenic amines exhibited a sustained decline from day 2 to day 8 and gradually recovered in the later phases of infection (Figure 1H, cluster 2). This affected almost exclusively essential- and semi-essential proteinogenic amino acids (His, Ile, Leu, Met, Trp, Val, Arg) (Figure 1H) and coincided with transcriptional regulation of amino acid-related metabolic pathways on day 8 (Figure 1F, 1G, 1H, S1F and Table S1C). Together, our integrated transcriptomic and proteomic analysis linked changes in the liver with altered systemic metabolite levels during chronic viral infection.

### Reduced food intake during viral infection mildly affects gene expression in the liver

Mice infected with LCMV strain clone 13 develop infection induced cachexia (Baazim et al., 2019) and display reduced food intake (Figure S2A). We therefore performed a pair-feeding experiment to investigate the impact of infection-associated changes in food to the metabolic reprogramming of the liver 8 days after infection. Uninfected mice received restricted amounts of food equivalent to the amounts consumed by infected mice. Subsequently, we collected liver tissue of infected, pair-fed and naïve animals and performed transcriptome analyses. These results indicated that most of the observed transcriptional changes were linked to viral infection and independent of anorexic behavior (Figure 2A, S2B and S2C). Uninfected pair-fed mice showed only minor differential gene expression changes that pertained mainly to lipid metabolism (Figure 2B, 2C and Table S4). Together, these experiments demonstrated that the majority of the observed virus-induced metabolic changes in the liver, including amino acid-related pathways, correlated with the infection status of the mice and were independent of altered food intake (Figure 1H and 2D).

**Figure 2:**
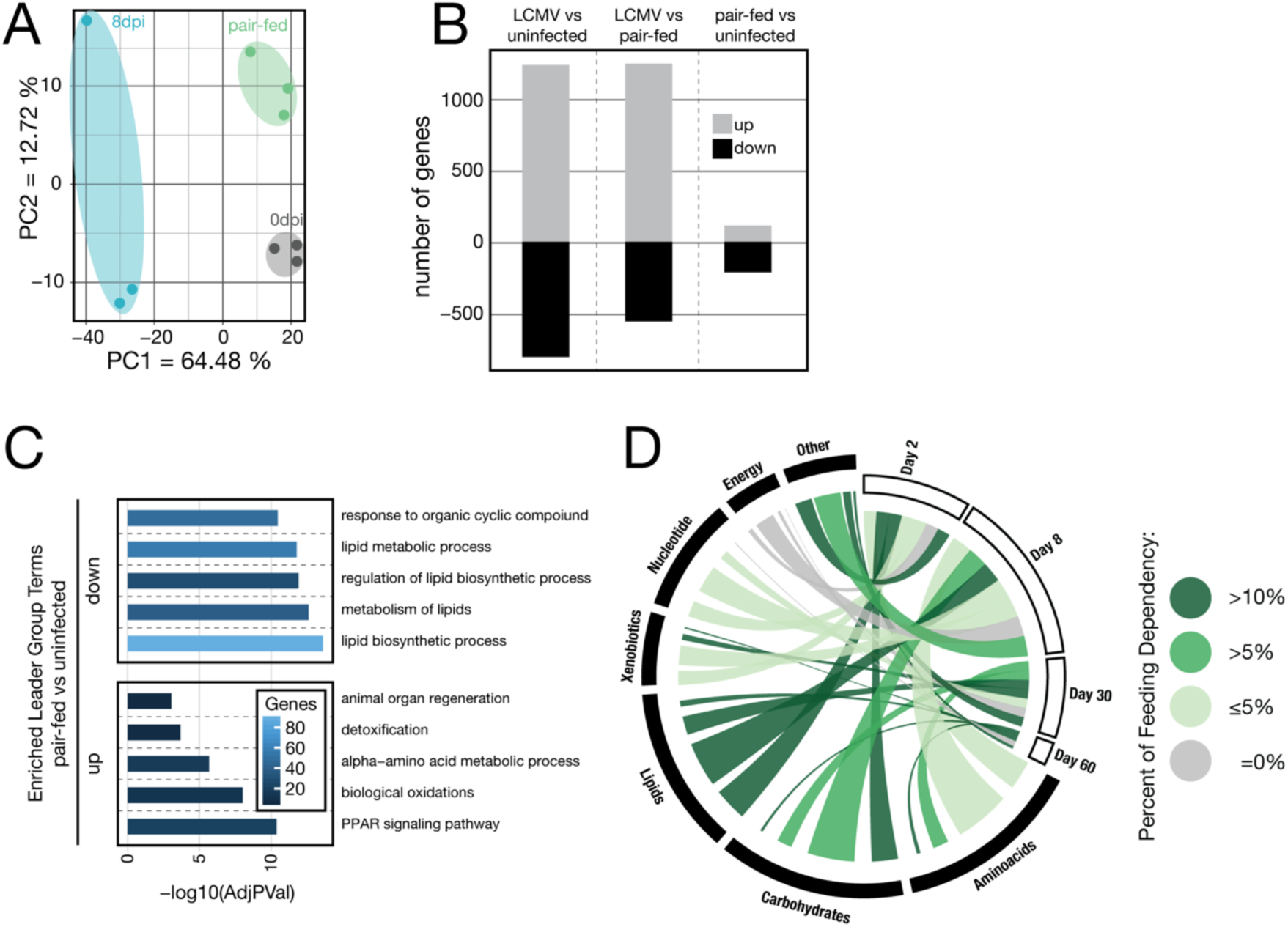
Reduced food intake upon LCMV infection mainly affects lipid metabolism associated genes in the liver. (A) Principal component analyses (PCA) of liver transcriptomes of naïve (0dpi), LCMV-infected (8dpi) and pair-fed (8 days) mice. (B) Significantly deregulated genes in liver tissue upon LCMV compared to naïve and pair-fed and between pair-fed and naïve animals. (C) Enriched GO terms and pathways (ClueGO) of significantly up- and downregulated genes of pair-fed versus naïve mice. (D) Metabolic genes in the liver (based on Figure 1G) affected by reduced food intake during LCMV infection.

### IFN-I signaling reprograms hepatic metabolism

The observed changes of hepatic amino acid metabolism became noticeable as early as 2 days, which coincides with peak serum levels of IFN-*α* and IFN-*β* in mice infected with LCMV (Bhattacharya et al., 2015). We, thus, aimed to dissect a potential role of early type I interferon signaling in the observed changes in the liver. To explore the potential impact of IFN-I signaling on liver parenchyma, we performed an extracellular metabolic flux analysis of primary murine hepatocytes upon stimulation with IFN-*β in vitro*. IFN-*β* affected proxies for mitochondrial respiration and glycolysis in Ifnar1-dependent way, (Figure 3A and 3B), indicating that IFN-I signaling affects metabolism of hepatocytes. To deconvolute tissue heterogeneity *in vivo* and assess the contribution of IFN-I signaling to the metabolic reprogramming in the liver, we took advantage of a genetic model of hepatocyte-specific ablation of *Ifnar1* (*Alb-Cre ERT2 Ifnar1^fl/fl^*, *Ifnar1^Δ/Δ^*) and respective littermate controls (*Alb-Cre ERT2 Ifnar1^+/+^*, *Ifnar1^+/+^*). These mice did not reveal any genotype-specific differences in viremia or serum concentration of IFN-*α* 1.5 days after infection (Figure S3A-C), suggesting that any potential differences seen in expression profiling of liver tissue from *Ifnar1^Δ/Δ^* vs. *Ifnar1^+/+^* mice is unlikely to be due to altered viral loads and/or systemic IFN-I responses. Next, we analyzed transcriptomic changes of liver tissue taken from uninfected and infected mice of either genotype at the peak and employed a limma (2×2 factorial) interaction model. This resulted in a set of 526 hepatocyte-intrinsic *Ifnar1*-regulated genes which were found to be associated with both classical ISG responses as well as metabolic processes (Figure 3C, 3D, S3D and Table S5A). The regulated genes could be divided into two major classes – *Ifnar1* stimulated (cluster I) and *Ifnar1* repressed (cluster II) genes (Figure 3C and Table S5B). The majority of induced genes were well-known classical ISGs encoding for antiviral effectors (cluster I, Figure 3D and Table S5C) (Schoggins et al., 2011). Interferon-repressed genes (IRGs), which are not that well characterized (Mostafavi et al., 2016; Schoggins et al., 2011), were found to be strongly enriched for metabolism-associated processes (cluster II, Figure 3D and Table S5C). In addition, clusters III and IV contained genes whose maintained expression depended on intact *Ifnar1* signaling and were also associated with metabolic processes (Figure 3D and Table S5C). Hepatocyte-intrinsic IFNAR1 signaling mainly regulated metabolic genes on day 2 after infection. Notably, a bioinformatic intersection with our longitudinal data obtained from chronically infected WT mice suggested that the IFN-I dependent regulation of many of these genes is maintained beyond these early time point (Figure 3E). Specifically, virus-induced gene regulation of amino acid-related pathways (Figure 1) was driven by hepatocyte-intrinsic IFNAR1 signaling (Figure 3E) and corresponded with the differentially regulated metabolites observed in infected WT mice (Figure 1H). To investigate whether these metabolic changes depended on hepatocyte-intrinsic IFNAR1 signaling, we performed metabolomics and found that systemic serum levels were indeed regulated by local IFNAR1 signaling of hepatocytes (Figure S3E). In a more stringent analysis using the limma (2×2 factorial) interaction model, we found 15 serum metabolites to be regulated by hepatocyte-intrinsic IFNAR1 signaling, including the semi-essential amino acid arginine and its downstream metabolite ornithine (Figure 3F). In summary, our data indicate that hepatocyte-intrinsic IFNAR1 acts as a transcriptional regulator of liver metabolism and results in changes of circulating metabolites.

**Figure 3:**
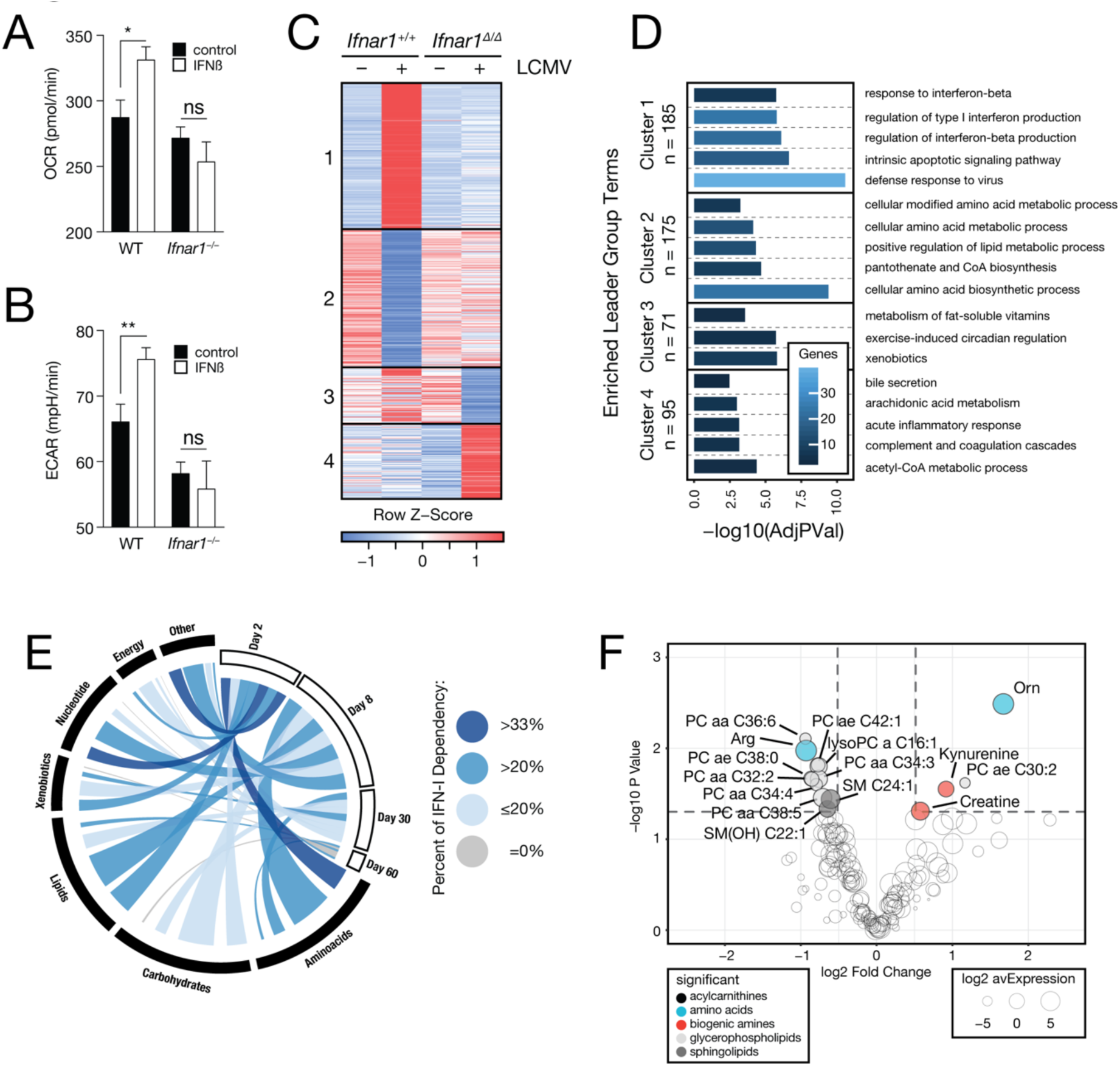
Hepatocyte-intrinsic *Ifnar1* signaling is a transcriptional regulator of liver metabolism and shapes systemic metabolism. (A) Oxygen consumption rate (OCR) and (B) extracellular acidification rate (ECAR) of wild type (WT) and *Ifnar1^−/−^* primary hepatocytes treated for 4h with IFN-*β* (n = 11). (C) Clustering by profile (FPKM, k-means, Pearson’s correlation) of genes significantly regulated by hepatocyte-intrinsic *Ifnar1* signaling (limma interaction model). (D) Enriched GO terms and pathways (ClueGO) of the identified clusters. (E) Metabolic genes in the liver (based on Figure 1G) affected by *Ifnar1* signaling in hepatocytes during LCMV infection. (F) Metabolites significantly regulated by hepatocyte-intrinsic *Ifnar1* signaling in naïve and LCMV-infected animals (limma interaction model). Symbols represent the arithmetic mean ±S.E.M. ns = not significant * P < 0.05 ** P < 0.01 (Student’s t-test).

### Hepatocyte-intrinsic *Ifnar1* signaling breaks the urea cycle

Arginine and ornithine are both key metabolites of the urea cycle. The involved enzymes are encoded by *Arg1*, *Otc* and *Ass1*, which are primarily expressed in the liver, as well as *Asl*, which is ubiquitously expressed (Uhlen et al., 2015). Expression analysis of liver tissue of uninfected and infected *Ifnar1^Δ/Δ^* and *Ifnar1^+/+^* mice highlighted that expression of *Otc* and *Ass1* was repressed in a hepatocyte-intrinsic IFNAR1-dependent manner (Figure 4A). *Arg1* was induced upon infection but this occurred independently of hepatocyte-intrinsic IFNAR1 signaling. In line with the IFNAR1-dependent increase of systemic ornithine levels upon infection (Figure 4A), these data indicate that IFNAR1 represses the degradation of ornithine via OTC in hepatocytes. The downregulation of *Asl* expression, together with decreased argininosuccinate, is expected to aggravate the infection-induced break in the urea cycle by limiting arginine resynthesis.

**Figure 4:**
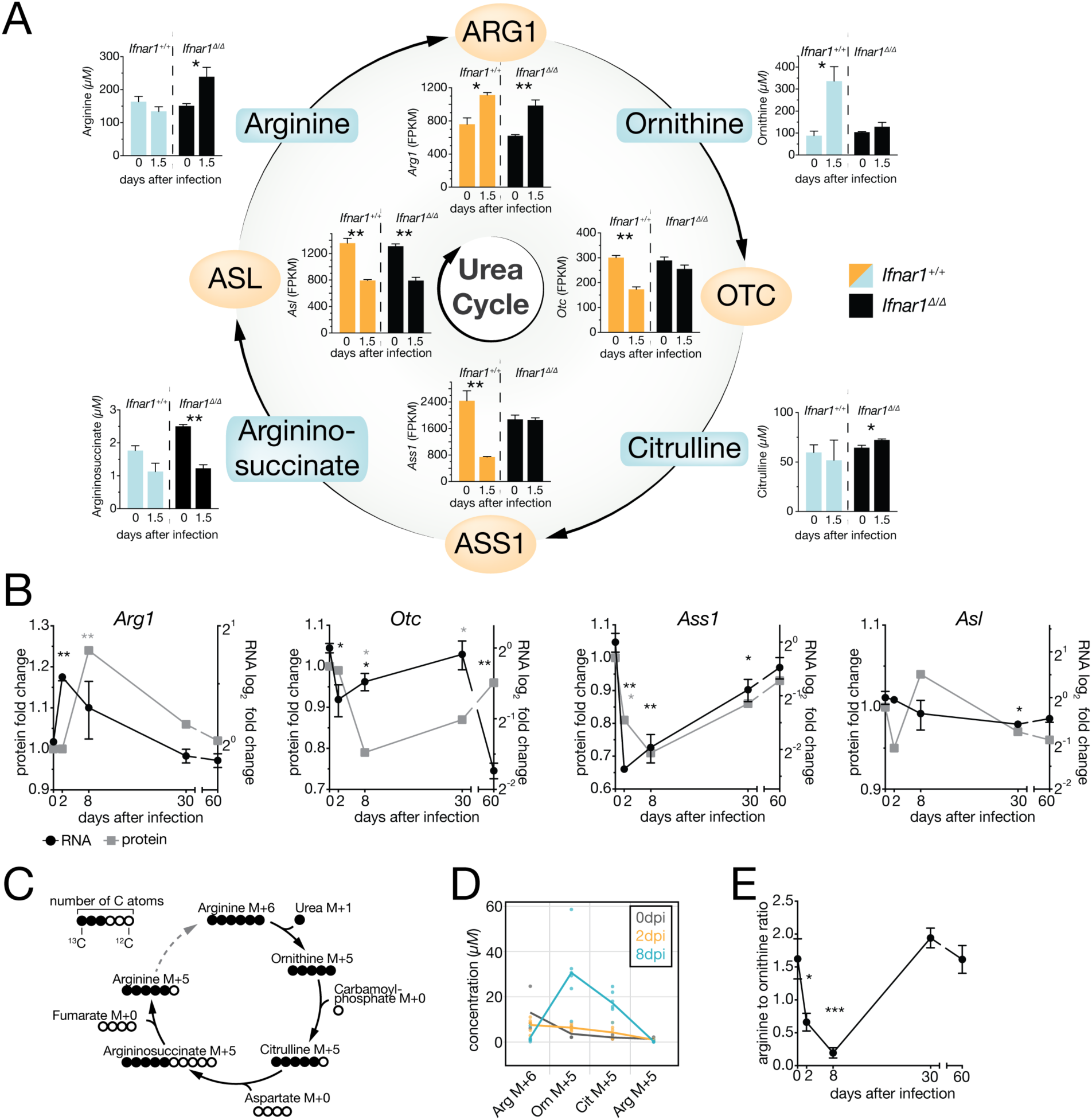
*Ifnar1* contributes to metabolic reprogramming of the urea cycle in hepatocytes to reduce the systemic arginine to ornithine ratio. (A) Depiction of the urea cycle, expression and concentrations of the associated genes and serum metabolites in naïve and LCMV clone 13 infected *Alb-Cre ERT2 Ifnar1^fl/fl^* (*Ifnar1^Δ/Δ^*) and *Ifnar1^+/+^* mice (n = 3). (B) Urea cycle gene transcript and protein abundances in liver tissue upon LCMV infection (n = 3). (C) Schematic depiction of the carbon flow in urea cycle starting from ^13^C_6_ arginine illustrating ^13^C labelled (full circles) and unlabeled ^12^C atoms of the respective metabolites. (D) Concentration of ^13^C labelled urea cycle metabolites at 0, 2 and 8 days after LCMV clone 13 infection (n = 3-6). (E) Systemic arginine to ornithine ratio of LCMV-infected C57BL/6J (wild type) animals (n = 8). Symbols represent the arithmetic mean ±S.E.M (B, E) or individual values (D). ns = not significant * P < 0.05 ** P < 0.01 *** P < 0.001 (Student’s t-test).

To test whether similar responses in the liver are seen in an inflammatory context and independent of LCMV infection. We administered the synthetic viral RNA analog and TLR3 agonist polyinosinic-polycytidylic acid (poly(I:C)) intraperitoneally to WT mice and collected liver tissue for gene expression analysis by real-time PCR. This revealed a poly(I:C)-induced downregulation of *Otc* and *Ass1* in the liver, corroborating the transcriptional regulation previously identified in our infection model (Figure S4A). These transcriptional changes of genes associated with the urea cycle were corroborated by the long-term transcript and protein kinetics of *Otc*, *Ass1* and *Arg1* in liver tissue of LCMV-infected mice, which were most pronounced on day 2 and 8 after infection and retained up to 30 days after infection, (Figure 4B). Of note, the decreased expression of *Otc* and Ass1 was independent of food intake (Figure S4B).

*In vivo* tracing of ^13^C_6_ heavy isotope labelled arginine revealed that viral infection increased ^13^C_6_ arginine degradation, consistent with the induction of *Arg1* (Figure 4C and 4D). Repression of *Otc* and *Ass1*, resulted in the accumulation of ^13^C_5_ ornithine and ^13^C_5_ citrulline, respectively (Figure 4D). Together with the reduction of ^13^C_5_ arginine, the end product of this pathway, these results support to the notion that viral infection initiates a break of the urea cycle as early as 2 days that persists at least until 8 day after infection (Figure S4C). Consistently, we observed a significant decrease of the arginine to ornithine ratio in the serum of infected mice (Figure 4E) as a result of both reduced arginine and increased ornithine levels (Figure S4D). Thus, hepatocyte-intrinsic IFN-I signaling reprograms the hepatic urea cycle and modulates the systemic arginine to ornithine ratio.

### Systemic arginine-ornithine homeostasis is a regulator of antiviral adaptive immunity

Since activated T cells are auxotrophic for arginine (Geiger et al., 2016; Murray, 2015), we hypothesized that an altered arginine to ornithine serum ratio might exert immunomodulatory function. To test this *in vitro*, primary naïve murine splenic CD8 T cells were activated with anti-CD3/CD28 antibodies for 3 days and IFN*γ* and TNF*α* was assessed by flow cytometry. CD8 T cells cultured in medium with 11.5 μM arginine, a concentration comparable to the observed changes *in vivo*, displayed reduced cytokine production compared to standard cell culture medium containing 1150 μM arginine (Figure 5A). Addition of ornithine, which is absent in standard cell culture medium, resulted in an additive suppressive effect on cytokine production (Figure 5A). This indicated that decreased concentrations of arginine impact CD8 T cell responses and that these effects are aggravated by a simultaneous increase of ornithine. Next, we aimed to experimentally uncouple the observed changes in the urea cycle from other co-occurring effects *in vivo* by treating WT mice with recombinant pegylated human Arginase 1 (recArg1) (Cheng et al., 2007). As expected, recArg1 converted arginine to ornithine in the serum and closely recapitulated the decreased arginine to ornithine ratio seen upon viral infection (Figure 5A and S5A). Administration of recArg1 did not affect the abundance of splenic T cells of infected and uninfected animals on day 8 after infection (Figure S5B). Yet, we recognized an impaired shift from naïve (CD62L^+^CD44^−^) to effector (CD62L^−^CD44^+^) T cells (Figure S5C and S5D). Further, the numbers of virus-specific CD8 T cells (Figure 5B) and their ability to produce the cytokines IFN*γ* and TNF*α* upon peptide re-stimulation (Figure 5C, 5D and 5E) were diminished upon recArg1 treatment. Virus-specific CD4 T cells were similarly affected (Figure S5E). CD8 T cell effector function remained impaired at least up to 50 days after infection (Figure 5F) and coincided with elevated viral loads in blood (Figure 5G) and organs (Figure S5F, S5G, S5H and S5I). Strikingly, treatment with recArg1 significantly ameliorated virus-induced CD8 T cell-mediated tissue damage as determined by reduced serum concentration of ALT and AST (Figure S5H and 5I). These data suggest that modulation of the systemic arginine to ornithine ratio regulates antiviral T cell responses and ameliorates virus-induced tissue damage.

**Figure 5:**
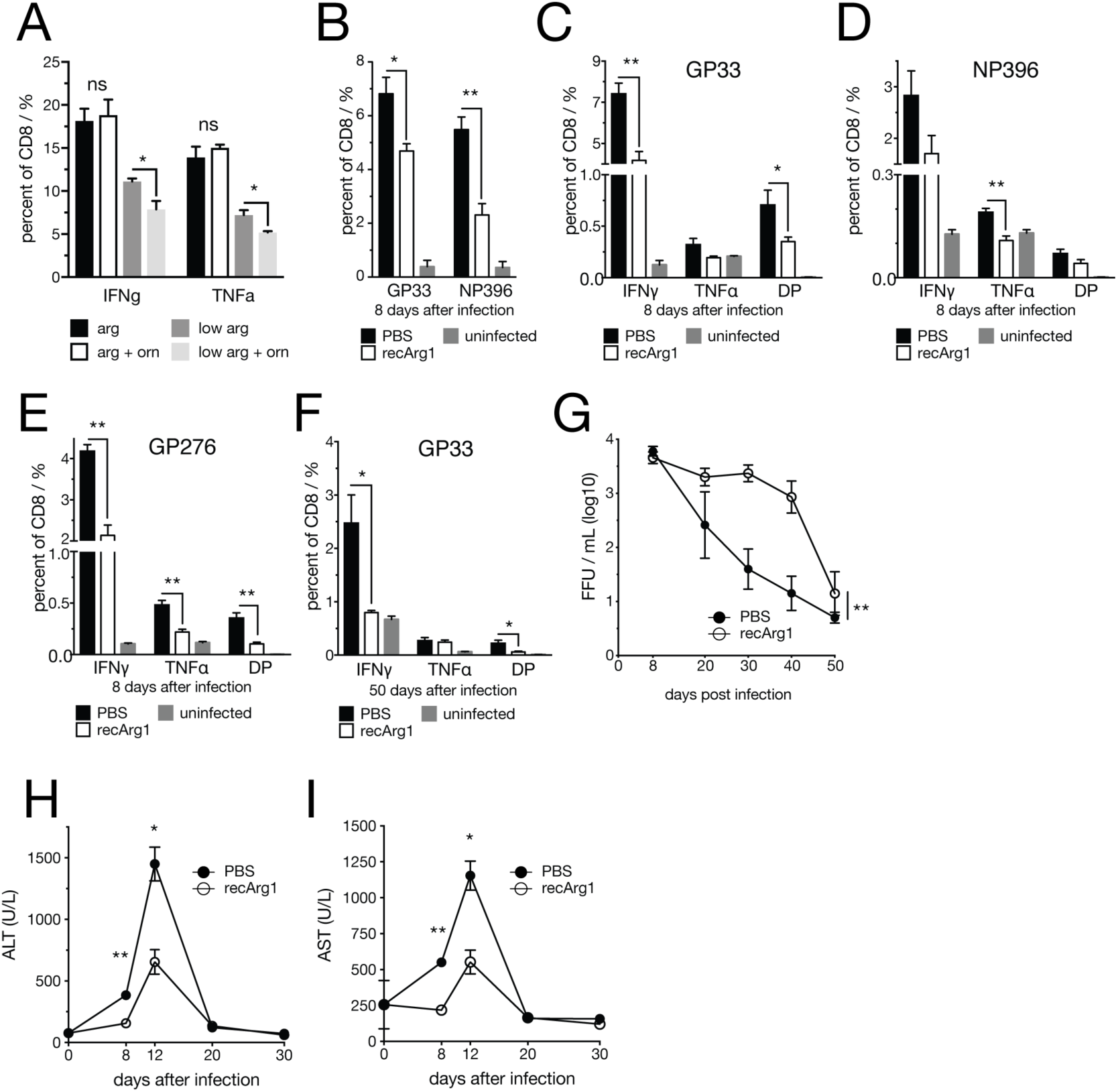
Arginase 1 treatment reduces antiviral CD8 T cell responses and ameliorates virus-induced hepatitis during LCMV Cl13 infection. (A) IFNγ and TNFα production of primary murine splenic CD8 T cells cultured in medium containing 1150µM (standard RPMI 1640 concentration) or 11.5µM L-arginine with or without 1150µM L-ornithine upon CD3/28 activation for 3 days. (B) GP33 and NP396-specific tetramer^+^ CD8 T cells. (C to E) IFNγ and TNFα production of virus-specific CD8 T cells upon recArg1 treatment 8 days after LCMV clone 13 infection. (F) IFNγ and TNFα producing GP33-specific CD8 T cells 50 days after infection. (G) Viral load in blood of upon recArg1 treatment (1 out of 3 representative experiments). (H) Serum ALT and (I) AST levels upon recArg1 treatment. n = 4-5. Symbols represent the arithmetic mean ±S.E.M. ns = not significant * P < 0.05 ** P < 0.01 (Student’s t-test (A to F, H, I) or two-way ANOVA (G)).

## Discussion

Metabolic adjustments are crucial for activation and differentiation of immune cells (Jha et al., 2015; Pearce and Pearce, 2013; Pearce et al., 2009). Our study identified an IFN-I-dependent mechanism whereby hepatocytes receive instructions to repress the transcription of genes with metabolic function. Interferon-repressed gene expression is still poorly understood in general. Yet, it is noticeable that IFN-I driven expression signatures from other organs and cell types do not appear to feature similar global repressive gene regulation (Mostafavi et al., 2016; Schoggins et al., 2011). This highlights that the liver can rapidly integrate inflammatory cues from the environment to modulate its intrinsic metabolism and, as a consequence, its systemic metabolic output to peripheral organs. This IFN-I induced metabolic reprogramming of the liver may give rise to an unfavorable environment for pathogens (Sanchez et al., 2018), optimize the metabolic milieu for mounting adaptive immune responses (Ma et al., 2017) or instruct tissue repair and regeneration to prevent extensive tissue pathology. None of these options are mutually exclusive, and host damage control may be achieved by modulating disease tolerance and/or reducing immune effectors (Medzhitov, 2008).

Next to other metabolic changes, we identified a regulatory effect of IFN-I on the urea cycle, a central metabolic pathway that is primarily expressed in the liver. Its primary function is the conversion of circulating toxic ammonia into non-toxic urea with a fumarate shunt connecting the urea cycle to the TCA cycle (Meijer et al., 1990). Fumarate can subsequently be metabolized to aspartate and α-ketoglutarate, two major precursors of *de novo* amino acid synthesis. Hence, the urea cycle is a central metabolic pathway that is crucial for both detoxication as well as cellular proliferation (Watford, 2003).

We modelled the altered levels of serum arginine and ornithine upon viral infection by administration of recombinant arginase and demonstrate a suppressive effect on virus-specific CD8 T cell responses. This is in line with nutrient availability as an important factor that may either boost or dampen effector functions of T cells (Chang et al., 2015; Geiger et al., 2016; Johnson et al., 2018; Ma et al., 2017; Sinclair et al., 2013; Van De Velde and Murray, 2016). A recent study identified that T cells may actively shape serum metabolite levels themselves, thereby creating potential functional feedback loops between systemic metabolism and T cell activation (Miyajima et al., 2017).

As a major hub of nitrogen metabolism, the urea cycle is not only linked to amino acid but also to polyamine and nucleotide metabolic pathways and may contribute to the formation of advanced liver disease (Li et al., 2019). Accordingly, urea cycle disorders (UCDs), gene deficiencies and the resulting changes in serum levels of the associated metabolites display severe health threats (Ah Mew et al., 1993). Downregulation or even loss of *Otc* and *Ass1* are common features of many cancers, limiting aspartate consumption by the urea cycle and instead feeding into *de novo* nucleotide synthesis associated with tumor growth (Feun et al., 2012; Lam et al., 2011; Lee et al., 2018; Rabinovich et al., 2015). *Otc* and *Ass1* are also crucial for *de novo* arginine synthesis. As a result, many tumors are auxotrophic for arginine, which renders arginine a metabolic vulnerability that is currently therapeutically exploited by recArg1 treatment of cancer patients (De Santo et al., 2018). In line, UCDs and in particular *Ass1*-deficiency of tumors potentially synergizes with checkpoint blockade treatment in tumors (Lee et al., 2018). Interestingly, reduced serum levels of arginine are associated with less liver tissue damage in chronic HBV infection while increased serum levels of arginine and arginine derivates have been associated with non-alcoholic fatty liver disease (Dumas et al., 2014; Li et al., 2011; Pallett et al., 2015; Soga et al., 2011). Finally, deregulated ornithine metabolism and increased synthesis of polyamines may also impact cellular proliferation as well as the replication of diverse groups of viruses (Casero et al., 2018; Mounce et al., 2016).

Here we demonstrate that virus-induced IFN-I signaling in hepatocytes leads to downregulation of *Otc* and *Ass1*. The simultaneous upregulation of *Arg1* during viral infection of the liver suggests that during infection, hepatocytes are reprogrammed towards increased arginine degradation and ornithine accumulation. In line with this, our ^13^C_6_ heavy isotope labeling experiments confirmed that viral infection increases the conversion of arginine to ornithine and leads to the accumulation of ornithine and citrulline. The observed infection-dependent reduction of the urea cycle end product ^13^C_5_ arginine further supported the notion of an Ifnar1-mediated break of the urea cycle. Previous studies also established a link between hepatic autophagy, hepatocyte-derived circulating ARG1 and the systemic arginine to ornithine ratio, leading to reduced tumor growth in mice (Poillet-Perez et al., 2018). These lines of evidence suggest that endogenous metabolic reprogramming of the urea cycle modulates the pathophysiology in different contexts of disease. Our results may also provide a potential explanation for why many non-hepatotropic infections result in concomitant hepatitis (Adams and Hubscher, 2006; Lalazar and Ilan, 2014). Taken together, our study provides evidence for an Ifnar1-dependent regulation of the central metabolic pathway of the hepatic urea cycle. This endogenous cytokine-driven mechanism of host-protection across organs could be targeted to ameliorate tissue pathology in infectious and inflammatory diseases (Figure S6).

## Acknowledgements

We want to thank Sarah Niggemeyer, Jenny Riede, Sabine Jungwirth and Patricia Schittenhelm for animal caretaking; Thomas Penz, Michael Schuster and Christoph Bock from the Biomedical Sequencing Facility at CeMM; Katja Parapatics from the Proteomics Facility at CeMM; and Jacques Colinge for data analyses. The following reagents were obtained through the NIH Tetramer Core Facility: LCMV MHCI tetramers GP33 (in PE), NP396 (in APC) and GP276 (in Alexa 488). This project was funded the European Research Council (ERC) under the European Union’s Horizon 2020 research and innovation program (grant agreement No 677006, “CMIL” to A.B.). A.L., A.Bh. and J.S.B. were supported by a DOC fellowship of the Austrian Academy of Sciences.

## Author Contributions

A. Lercher, A.P. and A. Bergthaler wrote the manuscript. A.L. and A.Bh. designed, performed *in vivo* and *in vitro* studies. M.F.S., L.K., T.P. performed *in vitro* and/or *in vivo* experiments. D.V., and K.L.B. contributed to proteomic analyses. A.L., A.P., M.C. and J.M. performed bioinformatic analyses. K.K., B.G., D.R. performed metabolite measurements. U.K. and P.N.C. provided reagents. H.B., G.S., J.S.B. and M.T. analyzed data and provided feedback. A.Be. supervised the study, designed and analyzed experiments.

## Competing interests statement

P.M. C. is the founder of Biocancer Treatment International Ltd. G.S. is scientific advisor of Biocancer Treatment International Ltd.

## Materials & Correspondence

Correspondence and requests for materials should be addressed to.

## Methods

### Mice

C57Bl/6J mice were originally obtained from The Jackson Laboratory, *Ifnar1*^fl/fl^ mice from Ulrich Kalinke (TWINCORE, Centre for Experimental and Clinical Infectiton Research, Hannover, Germany) and Cre-Alb ERT2 mice from Pierre Chambon (Institut de Génétique et de Biologie Moléculaire et Cellulaire, Illkirch, France). Mice were bred and maintained under specific pathogen-free conditions at the Institute for Molecular Biotechnology of the Austrian Academy of Sciences, Vienna, Austria. Animal experiments were conducted in individual ventilated cages according to the respective animal experiment licenses (BMWFW-66.009/0199-WF/V/3v/2015 and BMWFW-66.009/0361-WF/V/3b/2017) approved by the institutional ethical committees and the institutional guidelines at the Department for Biomedical Research of the Medical University of Vienna. Mice were age- and sex-matched within experiments. Male and female mice were used interchangeably between experiments and no striking sex-differences were observed.

### Pharmacological perturbations

Pegylated recombinant human Arginase 1(PEG-BCT-100, Bio-Cancer Treatment International Ltd.) was administered to mice via intraperitoneal injection at a dosage of 50 mg/kg. Mice were treated every third day. Treatment was started one day prior to LCMV infection. Control mice received the same volume of PBS.

### Metabolite tracing

Naïve or LCMV-infected C57Bl/6J mice were given an intravenous bolus of 500µg ^13^C_6_ labelled arginine (Cambrige Isotope Laboratories, CLM-2265-H-PK) dissolved in PBS. Mice were sacrificed 20 minutes afterwards and serum was harvested and stored at −80°C.

50 µL of serum were mixed with 450 µL of methanol and 250 µL of water and vortexed for 10 seconds. Next, 450 µL of chloroform were added, mixed by vortexing for 10 seconds, incubated on ice for 5 minutes and vortexted again for 10 seconds before centrifuging them fot 10 minutes at 1000 g. The upper aqueous phase was collected, dried (nitrogen evaporator) and reconstituted in 50 µL of methanol. Samples were centrifuged for 10 minutes at 1000 g and supernatants were used for LC-MS analysis.

A Vanquish UHPLC system (Thermo Scientific) coupled to an Orbitrap Fusion Lumos (Thermo Scientific) mass spectrometer was used for the LC-MS analysis. The chromatographic separation for samples was carried out on an ACQUITY UPLC BEH Amide, 1.7 µm, 2.1×100 mm analytical column (Waters) equipped with a VanGuard: BEH C18, 2.1×5mm pre-column (Waters). The column was maintained at a temperature of 40°C and 2 µL sample were injected per run. The mobile phase A was 0.15% v/v formic acid in water and mobile phase B was 0.15% v/v formic acid in 85% v/v acetonitrile with 10 mM ammonium formate. The gradient elution with a flow rate 0.4 mL/min was performed with a total analysis time of 17 min. The mass spectrometer was operated in a positive electrospray ionization mode: spray voltage 3.5 kV; sheath gas flow rate 60 arb; auxiliary gas flow rate 20 arb; capillary temperature 285°C. For the analysis a full MS scan mode with a scan range m/z 50 to 400, resolution 120000, AGC target 2e5 and a maximum injection time 50 ms was applied. The data processing was performed with the TraceFinder 4.1 software (Thermo Scientific).

### polyIC treatment

Mice were challenged with polyIC (invivogen, #tlrl-pic) via intraperitoneal injection at a dosage of 4 mg/kg. Control mice received PBS. Liver tissue was harvested and analyzed via RT-qPCR at the indicated time points.

### Isolation of peritoneal macrophages

Mice were sacrificed via cervical dislocation and the peritoneal cavity was flushed with 10 mL PBS. Cell pellets were resuspended and plated in RPMI medium (Gibco) containing 10% FCS (PAA) and 1% Penicillin-Streptomycin-Glutamine (Thermo Fisher Scientific). Medium was exchanged after 3 hours and cells were stimulated.

### Isolation of primary murine hepatocytes

Mice were anesthetized (Ketamine/Xylazine: 1:3, 0.1 ml/10 g mouse) and the liver was cannulated via the portal vein and perfused with 20 mL HBSS (Gibco) containing 0.5 mM EGTA (Sigma) followed by digestion of the liver with 20 mL L15 medium (Gibco) containing 40 mg/L Liberase (Roche) at a rate of 5 mL/min. The liver was isolated, placed in a petri dish with digestion medium (L15 with 40 mg/L Liberase) and the liver capsule was diligently removed. Cells were centrifuged at 5G for 5 min at 4°C, resuspended in William’s E medium containing 10% FCS (PAA) and 1% Penicillin-Streptomycin-Glutamine (Thermo Fisher Scientific) and plated. Cell were stimulated at the time of plating.

### Cytokine determination

IFNα serum levels were determined via ELISA as described previously (Bhattacharya et al., 2015). IFNγ in serum was measured by ELISA according to the manufacturer’s instructions (Mouse IFN-γ ELISA MAX™ Deluxe, Biolegend). For both assays, serum samples were pre-diluted 1:10 to 1:20.

### Viruses and infection

Lymphocytic choriomeningitis virus (LCMV) was grown on BHK-21 cells and titer was determined in a modified focus forming assay using Vero cells (Bhattacharya et al., 2015). Mice were intravenously infected with 2×10^6^ focus forming units (FFU) of LCMV.

### Blood chemistry

Mouse serum was pre-diluted 1:8 in PBS and levels of alanine aminotransferase (ALT) and aspartate aminotransferase (AST) were spectrophotometrically analyzed using a Cobas C311 Analyzer (Roche). Blood urea concentrations were determined using a colorimetric assay according to the manufacturer’s instructions (Sigma, MAK006).

### RNA isolation and real-time PCR

Tissues were homogenized using a TissueLyser II (Qiagen). Total RNA was extracted from homogenized liver tissue using QIAzol lysis reagent as per the manufacturer’s instructions (79306, Qiagen). Reverse transcription from RNA to cDNA was carried out using random primers and the First Strand cDNA Synthesis Kit (K1612, Thermo Fisher Scientific). Real-time PCR was performed with Taqman Fast Universal PCR Mastermix (4352042, Thermo Fisher Scientific) and Taqman Gene Expression Assays (Thermo Fisher Scientific) for *Otc* (Mm00493267_m1) and *Ass1* (Mm00711256_m1). Expression levels of LCMV NP and EF1a were measured by corresponding probe and primer sets as described previously (Gilchrist et al, 2006, Pinschewer et al, 2010).

### Metabolic flux measurements

Oxygen consumption rate (OCR) and extracellular acidification rate (ECAR) were determined on a Seahorse XFe96 Analyzer (Agilent) using the Seahorse XF Cell Mito Stress test kit (Agilent, 103015-100). 150.000 peritoneal macrophages and 25.000 primary hepatocytes were plated per well respectively. In brief, cells were treated with selected stimuli for the indicated time points. Prior to measurement, media was changed to XF Base Medium (Agilent 102353-100) containing glucose (10 mM), sodium pyruvate (1 mM) and L-glutamine (2 mM) and cells were incubated for 1h. The assay was run according to the manufacturer’s instructions. Oligomycin (2 µM primary hepatocytes, 1 µM peritoneal macrophages), Carbonyl cyanide-p-trifluoromethoxyphenylhydrazone (FCCP, 0.25µM primary hepatocytes, 1 µM peritoneal macrophages) and Rotenone/Antimycin A (500 nM) were subsequently injected into wells at the desired time points. Raw data was analyzed using Wave Desktop Software (Agilent, version 2.0) and exported and graphed in GraphPad Prism (GraphPad Sorftware, version 7.0a).

### CD8 T cell isolation and *in vitro* activation

Spleens of mice were harvested and dissociated through a 40µm cell strainer (Falcon). The pellet was resuspended in 1mL red blood cell lysis buffer (Invitrogen, #00-4970-03) and incubated at room temperature for 1 minute. Subsequently the reaction was stopped by addition of 9mL PBS. Cells were counted and CD8 T cells isolated using a magnetic activated cell sorting negative selection (MACS) kit according to the manufacturer’s instructions (Miltenyi Biotec, #130-104-075). 96 well plates were coated over night at 4°C with 1µg/mL anti CD3e (BD, #553238) and 2µg/mL anti CD28 (BD, #553295) in a total volume of 60µL PBS per well. Wells were washed with PBS and 5×10^4^ cells were plated per well in RPMI 1640 medium for SILAC (Thermo Fisher Scientific, #88365), supplemented with 10% dialyzed FCS (Thermo Fisher Scientific, # A3382001), 1% Penicillin-Streptomycin-Glutamine (Thermo Fisher Scientific), 50µM β-mercaptoethanol (Sigma, #M6250) and 20 U/mL IL2 (Thermo Fisher Scientific, #34-8021-82). Medium was supplemented with indicated concentrations of L-arginine (Sigma, #A5006) and/or L-ornithine (Sigma, #O2375) and cultured up to 72h before proceeding with intracellular cytokine staining.

### Flow cytometry

To sample blood, 3-4 drops of blood were collected in 1 mL MEM medium (Gibco) supplemented with 2000 U/L heparin (Ratiopharm). Red blood cells were lysed by adding 500 µL red blood cell lysis buffer and incubating at room temperature for 1 minute. Samples were spun down (1500 rpm), resuspended in 100 µL PBS and plated in a 96 well plate. Spleens were dissociated into a single cell suspension using a 40 µm cell strainer (Falcon) and resuspended in 10 mL of PBS (Gibco). An aliquot of cell suspension was used for counting to calculate total number of cells per spleen. 200 µL per sample (approx. 2×10^6^ cells) were plated in a 96 well plate. Plates were spun and supernatants were removed.

For tetramer staining, cells were resuspended in 25 µL PBS containing GP33 (1:500) and NP396 (1:250) tetramers (NIH Tetramer Core Facility) and incubated at 37°C for 15 min. Next, 25 µL PBS containing anti-CD16/32 (1:200, clone: 93) were added and incubated for 10 minutes at room temperature. 25 µL of a master mix of the desired surface marker antibodies (CD8.2b: Pacific Blue clone 53-5.8; CD4: FITC, clone H129.19; CD4: PE, clone GK1.5; CD3: APC, clone 145-2C11; CD44: BV605, clone IM7; CD62L: AF700, clone MEL-14; CD19: APC-Cy7, clone 6D5; all Biolegend; 1:200 in PBS) and eBioscience Fixable Viability Dye eFluor 780 (1:2000 in PBS) were added and cells were incubated for 20 minutes at 4°C. Cells were washed with FACS buffer (PBS, 2%FCS) and fixed in 4% Paraformaldehyde (Sigma) in PBS for 10 minutes. Subsequently samples were washed twice with FACS buffer, resuspended in 100 µL and analyzed by flow cytometry.

For intracellular cytokine staining (ICS), cell pellets were resuspended in 50 µL of RPMI 1640 medium (Gibco) supplemented with 10% FCS (PAA) and 1% Penicillin-Streptomycin-Glutamine (Thermo Fisher Scientific), 50µM β-mercaptoethanol (Sigma), containing LCMV peptides (1:1000, Peptide 2.0 Inc.) and Protein Transport Inhibitor Cocktail (1:500, Thermo Fisher Scientific). Cells were incubated for 4 hours at 37°C and then surface antigens were stained as described above. Afterwards, a master mix of desired antibodies in 25 µL FACS buffer containing 0.05% saponin (Sigma) against intracellular antigens of interest were added and incubated for 90 minutes at 4°C (IFNγ: PE-Cy7, clone XMG1.2; IL2: PE, clone JES6-5H4; TNFα: APC, clone MP6-XT22; all Biolegend. all 1:200). Next, cells were washed twice with FACS buffer, resuspended in 100 µL and analyzed by flow cytometry.

### RNAseq

RNA quality and integrity was assessed via an Experion RNA HighSense chip (Biorad). The library for RNAseq was prepared using the TruSeq RNA sample preparation kit v2 (Illumina) according to the manufacturer’s protocol. Quality control analysis was performed on all samples of the cDNA library by Experion DNA Analysis chip (Biorad) and Qubit Fluorometric quantitation (Life Tech). 7 or up to 17 samples were multiplexed per lane and run on a 50bp single-end flow cell in a HiSeq2000 or HiSeq3000 sequencer (Illumina), respectively. Called bases by the Illumina Realtime Analysis software were converted into BAM format using Illumina2bam and demultiplexed using BamIndexDecoder (https://github.com/wtsi-npg/illumina2bam). The RNAseq analysis pipeline was performed with Tuxedo. Reads were mapped on the mouse reference genome (Mus musculus, Ensembl e87, December 2016) using TopHat2 (v2.0.10). Cufflinks (v2.2.1) was employed to assemble transcripts from spliced read alignments, using the Ensembl e87 transcriptome as the reference as well as *de novo* assembly of transcript models. Further, differential analysis of gene expression was quantified with Cuffdiff (v2.2.1). Transcriptome sets of all replicates for each sample group were combined with Cuffmerge. Expression values in graphs are reported as FPKM (fragments per kilobase pf transcript per million). Differential gene expression is attested based on expression level *≥* 1 FPKM, adjusted p value *≤* 0.05 and absolute log2 fold-change of 1 (heatmap) or 0.6 (circos plot).

### Principal component analysis

Principal component analysis (PCA) was performed on the gene set with a minimum average expression level across conditions of 5 FPKM. Only the 10% most variable (computed on the coefficient of variance) genes were considered for the PCA analysis.

### Interaction model

For the interaction model (2×2 factorial design) of naïve versus LCMV-infected *Ifnar1^Δ/Δ^* and *Ifnar1^+/+^*, we quantified gene expression as the number of reads covering each gene. The gene expression on the mouse Ensembl e87 transcripts was quantified from the previously TopHat2 mapped reads with featureCounts (Liao et al., 2014). Raw read counts were normalized with the voom (Law et al., 2014) function of the limma package(Smyth, 2004). Normalized expression values, reported as log2 counts per million (CPM), were then processed through limma’s empirical Bayes models. We implemented limma’s interaction model as a two (*Ifnar1^+/+^*, *Ifnar1^Δ/Δ^*) by two (naïve, LCMV-infected) factorial design. Genes differentially modulated in the interaction model have been selected based on a minimum log2 CPM of 0, an adjusted p value *≤* 0.05 and a minimum log2 fold-change absolute value of 1 (heatmap) or 0.6 (circos plot).

### Hierarchical clustering

Hierarchical clustering of different gene sets FPKM or CPM expression as well as metabolite abundance values were performed using a Pearson distance measure with an average clustering method. A k-means ++ (z-norm) clustering approach using ExpressCluster software v1.3 (http://cbdm.hms.harvard.edu/LabMembersPges/SD.html) was performed on the union of differentially modulated genes or metabolites.

### Quantitative proteomics

Three TMT 6-plex runs were carried out to monitor the changes in liver protein abundance during the entire course of infection. The first run included biological replicates from day 2 and day 8 after infection along with the uninfected controls. The second run included replicate samples of uninfected controls, day 30 and day 60 after infection. To account for the effect of aging, the third run included uninfected controls at day 0 and day 123, along with infected samples from day 123 (all in biological replicates). Identical uninfected controls (day 0) were included in all three TMT 6-plex runs as an internal control to monitor reproducibility between the runs. Liver tissues were homogenized (TissueLyser II, Qiagen) in 1.5 mL 50 mM HEPES buffer, pH 8.5 supplemented with 2% sodium dodecyl sulphate (SDS). The protein concentrations were determined by the bicinchoninic acid assay (BCA, Pierce Biotechnology, Thermo Scientific, IL). Further processing was adapted from a filter-aided sample preparation (FASP) method previously described (Manza *et al*, 2005, Wisniewski *et al*, 2009). For each sample, 100 µg of the liver tissue lysate was reduced with 100 mM dithiothreitol (DTT) and transferred into VIVACON 500 filter unit (Vivaproucts Inc., Littleton, MA). SDS-containing buffer was removed from the sample by centrifugation and exchanged with 8 M urea in 100 mM Tris-HCl buffer. Proteins were alkylated with 50 mM iodoacetamide and washed with 50 mM triethyl ammonium bicarbonate (TEAB). Finally, porcine trypsin (Promega Corp., Madison, WI) was used for protein digestion in an enzyme to protein ratio of 1:100 w/w.

For relative protein quantitation, six samples from each run were separately derivatized with TMT 6-plex reagents (ABI, Framingham, MA) according to the instructions provided by the manufacturer. The combination of the TMT 6-plex labels was as follows: (i) Run 1: day 0 uninfected (TMT 126 and 127), day 2 infected (TMT 128 and 129) and day 8 infected (TMT 130 and 131); (ii) Run 2: day 0 uninfected (126 and 127), day 30 infected (128 and 129) and day 60 infected (130 and 131); and (iii) Run 3: day 0 (126 and 127), day 123 uninfected (128 and 129) and day 123 infected (130 and 131).

The TMT-labelled tryptic digests were pooled and concentrated by solid phase extraction (SPE) (MacroSpin columns 30-300 µg capacity, The Nest Group Inc., Southborough, MA, USA). Samples were treated with 20 mM ammonium formate prior to injection onto a Phenomenex column (150×2.0 mm Gemini®NX-C18 3 µm 110Å; Phenomenex, Torrance, CA, USA) in an Agilent 1200 series HPLC (Agilent Biotechnologies, Palo Alto, CA) with UV detection at 214 nm. HPLC solvent A consisted of 20 mM ammonium formate, pH 10 in 5% acetonitrile. Peptides were separated at flow rate of 100 µL/min and eluted from the column with a non-linear gradient ranging from 0 to 100% solvent B (20 mM ammonium formate, pH 10 in 90% acetonitrile). Seventy-two time-based fractions were collected, acidified and pooled into 50 HPLC vials based on the UV trace. After removal of organic solvent in a vacuum centrifuge, samples were reconstituted to 10 µL with 5% formic acid (Bennett et al, 2011). Individual fractions were further analyzed at pH 2.4 by Agilent 1200 nano-HPLC system (Agilent Biotechnologies, Palo Alto, CA) coupled to a hybrid LTQ Orbitrap Velos mass spectrometer (ThermoFisher Scientific, Waltham, MA). Data was acquired utilizing the Xcalibur software version 2.1. Briefly, single fractions were loaded onto a trap column (Zorbax 300SB-C18 5 µm, 5 × 0.3 mm, Agilent Biotechnologies, Palo Alto, CA) with a binary pump at a flow rate of 45 µL/min. Solvents for LCMS separation were composed of 0.1% trifluoracetic acid (TFA) in water (solvent A) and 0.1% TFA in 70% methanol and 20% isopropanol (solvent B). The peptides were eluted by back-flushing from the trap column onto a 16 cm fused silica analytical column with an inner diameter of 50 µm packed with C18 reversed-phase material (ReproSil-Pur 120 C18-AQ, 3 µm, Dr. Maisch GmbH, Ammerbuch-Entringen, Germany). Elution was achieved with a 27 min gradient ranging from 3 to 30% solvent B, followed by a 25 min gradient from 30 to 70% solvent B and, finally, a 7 min gradient from 70 to 100% solvent B at a constant flow rate of 100 nL/min. The analysis was performed in a data-dependent acquisition mode. The 10 most intense ions were isolated and fragmented by high-energy collision-induced dissociation (HCD) for peptide identification and relative quantitation of TMT reporter ions. Dynamic exclusion for selected ions was 60 s and a single lock mass at *m/z* 445.120024 (Si(CH3)2O)6)20 (Olsen et al, 2005) was used for internal mass calibration with the target loss mass abundance of 0%. Maximal ion accumulation time allowed was 500 ms and overfilling of the C-trap was prevented by automatic gain control set to 10^6^ ions for a full FTMS scan and 5×10^5^ ions for MS^n^ HCD. Intact peptides were detected in the Orbitrap mass analyser at resolution of 30,000 with the signal threshold of 2,000 counts for triggering an MSMS event. The maximum ion scan time was set to 200 ms for acquiring 1 microscan at a resolution of 7500.

### Mass spectrometry data processing

The acquired raw MS data files were processed with msconvert (ProteoWizard Library v2.1.2708) and converted into MASCOT generic format (mgf) files. Peptides were identified by searching the resultant peak lists against the SwissProt mouse database version v2013.01_20130110 (24615 sequences; 14280050 residues) with the search engines MASCOT (v2.3.02, MatrixScience, London, UK) and Phenyx (v2.5.14, GeneBio, Geneva, Switzerland). Submission to the search engines was done via a Perl script that performs an initial search with relatively broad mass tolerances (MASCOT only) on both the precursor and fragment ions (±10 ppm and ±0.6 Da, respectively). High-confidence peptide identifications were used to recalibrate all precursors and fragment ion masses prior to a second search with narrower mass tolerances (±4 ppm and ±0.025 Da). Trypsin was chosen as cleavage specificity with the maximum of 1 miscleavage site allowed. Carbamidomethyl cysteine, N-terminal and lysine-modified TMT 6-plex were set as fixed modifications, whereas oxidized methionine was set as a variable modification.

To validate the proteins, MASCOT and Phenyx output files were processed by internally developed parsers. Proteins with ≥2 unique peptides above a score T1, or with a single peptide above a score T2, were selected as unambiguous identifications. Additional peptides for these validated proteins with score >T3 were also accepted. For MASCOT and Phenyx, T1, T2, and T3 peptide scores were equal to 16, 40, 10 and 5.5, 9.5, 3.5, respectively (P-value <10^−3^). The validated proteins retrieved by the two algorithms were merged, any spectral conflicts discarded, and grouped according to shared peptides. A false positive detection rate (FDR) of <1% and <1% was determined for proteins and peptides, respectively, by applying the same procedure against a reversed database.

The log2 fold-change cutoffs for differential protein modulation were determined based on pairwise comparisons of protein abundances between the two replicates of uninfected control samples across the 3 independent runs. The 2.5% and 97.5% quartiles of the inter-replica pairwise log2 fold-change was computed. Based on the quartile values, we have set the cutoff at 0.25 and −0.25 for up- and down-modulated proteins, respectively. Additional to the log2 fold-change, statistical significance of observed changes was calculated by Isobar software (Breitwieser et al., 2011). Adjusted P-value ratio as well as the P-value samples as calculated by Isobar were asked to be less than 0.05.

### Targeted LC-MS based metabolite quantification

Tissue samples were homogenized using a Precellys 24 tissue homogenizer (Precellys CK14 lysing kit, Bertin). Per mg tissue, 3µL of 80% (v/v) methanol were added. 10 µL of homogenized tissue sample or serum were mixed with 10 µL of an isotopically labeled internal standard mixture in a hydrophobic 96well filter plate. Aliquots of 300 µL of methanol were added and mixed for 20 min at 450 rpm. Afterwards, the sample extracts were collected in a 96-well plate by centrifuging the filter plate for 5 min at 500 g. A Vanquish UHPLC system (Thermo Scientific) coupled with an Orbitrap Q Exactive (Thermo Scientific) mass spectrometer was used for the LC-MS analysis. The chromatographic separation for samples was carried out on an ACQUITY UPLC BEH Amide, 1.7 µm, 2.1×100 mm analytical column (Waters) equipped with a VanGuard: BEH C18, 2.1×5mm pre-column (Waters). The column was maintained at a temperature of 40°C and 2 µL sample were injected per run. The mobile phase A was 0.15% formic acid (v/v) in water and mobile phase B was 0.15% formic acid (v/v) in 85% acetonitrile (v/v) with 10 mM ammonium formate. The gradient elution with a flow rate 0.4 mL/min was performed with a total analysis time of 17 min. The Orbitrap Q Exactive (Thermo Scinetific) mass spectrometer was operated in an electrospray ionization positive mode, spray voltage 3.5 kV, aux gas heater temperature 400°C, capillary temperature 350°C, aux gas flow rate 12. The metabolites of interest were analyzed using a full MS scan mode, scan range m/z 50 to 400, resolution 35000, AGC target 1e6, maximum IT 50ms. The Trace Finder 4.1 software (Thermo Scientific) was used for the data processing. Seven-point linear calibration curves with internal standardization and 1/x weighing was constructed for the quantification of metabolites.

Metabolite quantifications using the AbsoluteIDQ p180 Kit (Biocrates Life Science AG) were performed as described previously (St John-Williams et al., 2017). In brief, per mg of tissue, 6µL ethanol/phosphate buffer (85:15 v/v ethanol/10mM phosphate buffer) were added and the tissue was homogenized (TissueLyser II, Qiagen). Samples were centrifuged at 5000g for 5 minutes at 4°C and the supernatants were transferred to a fresh tube and used for analyses. For serum samples, blood was collected from mice in blood collection tubes and centrifuged at 12000g for 5 min to obtain sera. The serum was transferred into a fresh tube and stored at −80°C until analyses. The samples were analyzed on a Xevo TQ-MS (Waters) mass spectrometer using an Acquity UHPLC (Waters) system, operated with MassLynx V4.1 (Waters). Samples and additional blanks, calibration standards and quality controls were prepared according to the user manual. The sample measurement and data analysis were carried out according to the user manual. The experiments were validated with the supplied software MetIDQ, Version 5-4-8-DB100-Boron-2607 (Biocrates Life Sciences). All metabolite abundances inferior to the limit of detection (LOD) were replaced with a value equal to half the LOD. We eliminated all metabolites that did not have an average abundance superior to LOD in at least one condition. Modulation of metabolites was assessed through a t-test between conditions. The cutoffs for the modulation were obtained based on all the inter-replica pairwise comparisons of wild-type uninfected samples. The 2.5 and 97.5 quartiles of these inter-replica log fold-change were used to define the cutoffs. The cutoffs were −0.44 and 0.65 for the whole liver dynamics serum metabolite measurements. For serum measurements of *Ifnar1^Δ/Δ^* and *Ifnar1^+/+^* mice at 1.5 days post infection, based on the same approach, we have set the cutoffs at −0.73 and 0.4. Significance was inferred based on a p value inferior to 0.05. For the interaction model (2×2 factorial design) of naïve versus LCMV-infected *Ifnar1^Δ/Δ^* and *Ifnar1^+/+^*, we implemented limma’s interaction model as a two (*Ifnar1^+/+^*, *Ifnar1^Δ/Δ^*) by two (naïve, LCMV-infected) factorial design. Significantly modulated metabolites in the interaction model were identified by an absolute log2 fold change of 0.7 and a p value inferior to 0.05.

### Enrichments and pathway analyses

Enrichment analyses on the union of differential modulated entities (transcripts and/or proteins) specific clusters were done in Cytoscape ClueGO (Bindea et al., 2009) v2.3.3, based on GO (Biological Processes, Molecular Functions, Immune System Process), InterPro, KEGG, Reactome and Wiki Pathways. Terms were called enriched based on a maximum p-value of 0.05 and a minimum of 2% gene overlap. GO Term Fusion and grouping was applied. Enriched groups where farther ranked according to the group Bonferroni step-down adjusted p-value.

For metabolic pathway enrichment analyses, we took the union of differentially expressed genes (expression *≥* 1 FPKM, absolute log2 fold-change *≥* 0.7, adjusted p value *≤* 0.05) and proteins (absolute log2 fold-change *≥* 0.25 and adjusted ratio p value < 0.05 and sample p-value < 0.05) across all time points. We extracted metabolism-associated genes from KEGG(Kanehisa and Goto, 2000) metabolic pathways database. Only the pathways with q value of enrichment < 0.05 were considered. The enriched pathways are represented as a circos plot (Krzywinski et al., 2009), with the width of each ribbon in a given category representing the percentage of genes at each time point among all genes leading to enrichments in the respective category. The color gradient (from ligher to darker) represents the percentage of pathways in each category that were found as enriched at a specific time point.

### Statistical information

Data is presented as arithmetic mean ±S.E.M. Statistical significances were calculated using a Student’s t-test when comparing two groups or using two-way ANOVA with Bonferroni correction when comparing longitudinal changes. * P < 0.05 ** P < 0.01 *** P < 0.001.

### Data availability

Proteomic data (PXD011122) and transcriptomic data (GSE123688, which includes GSE118703, GSE123684 and GSE118819) are deposited in the PRIDE respectively GEO databases.

---

**Supplementary Figure S1:**
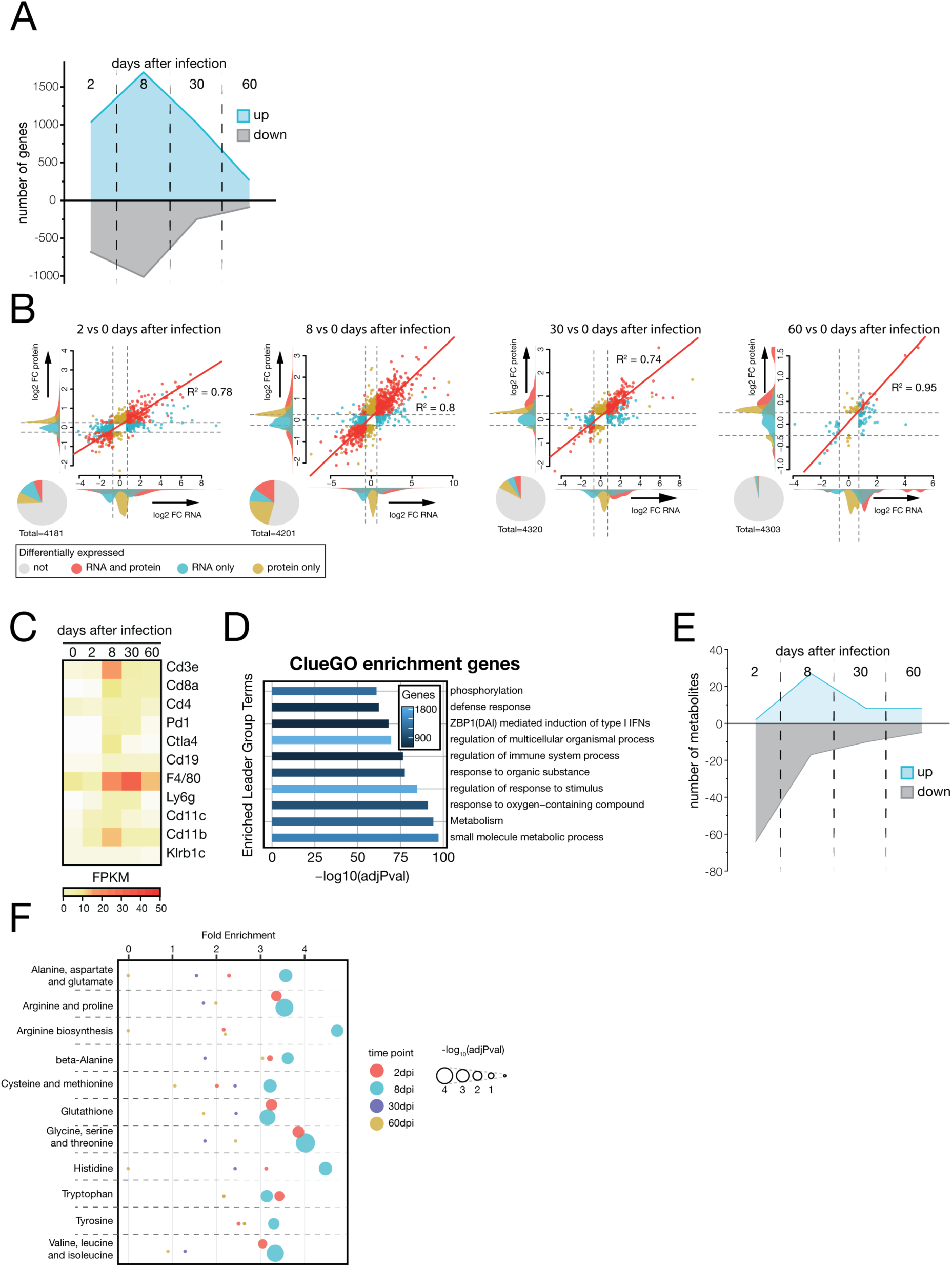
Systems biology approach to transcriptomic, proteomic and metabolomic changes in liver tissue and systemic metabolism during LCMV Cl13 infection. (A) Number of regulated genes in liver tissue upon LCMV infection. (B) Correlation of transcriptomic and proteomic changes at the indicated time points. (C) FPKM values of indicated immune cell markers in liver tissue. (D) Enriched GO terms and pathways (ClueGO) over the union of differentially expressed genes. (E) Number of up- or downregulated metabolites of total significant metabolites in serum and liver upon LCMV infection. (F) Enrichment analyses for amino acid metabolic pathways (KEGG) during infection.

**Supplementary Figure S2:**
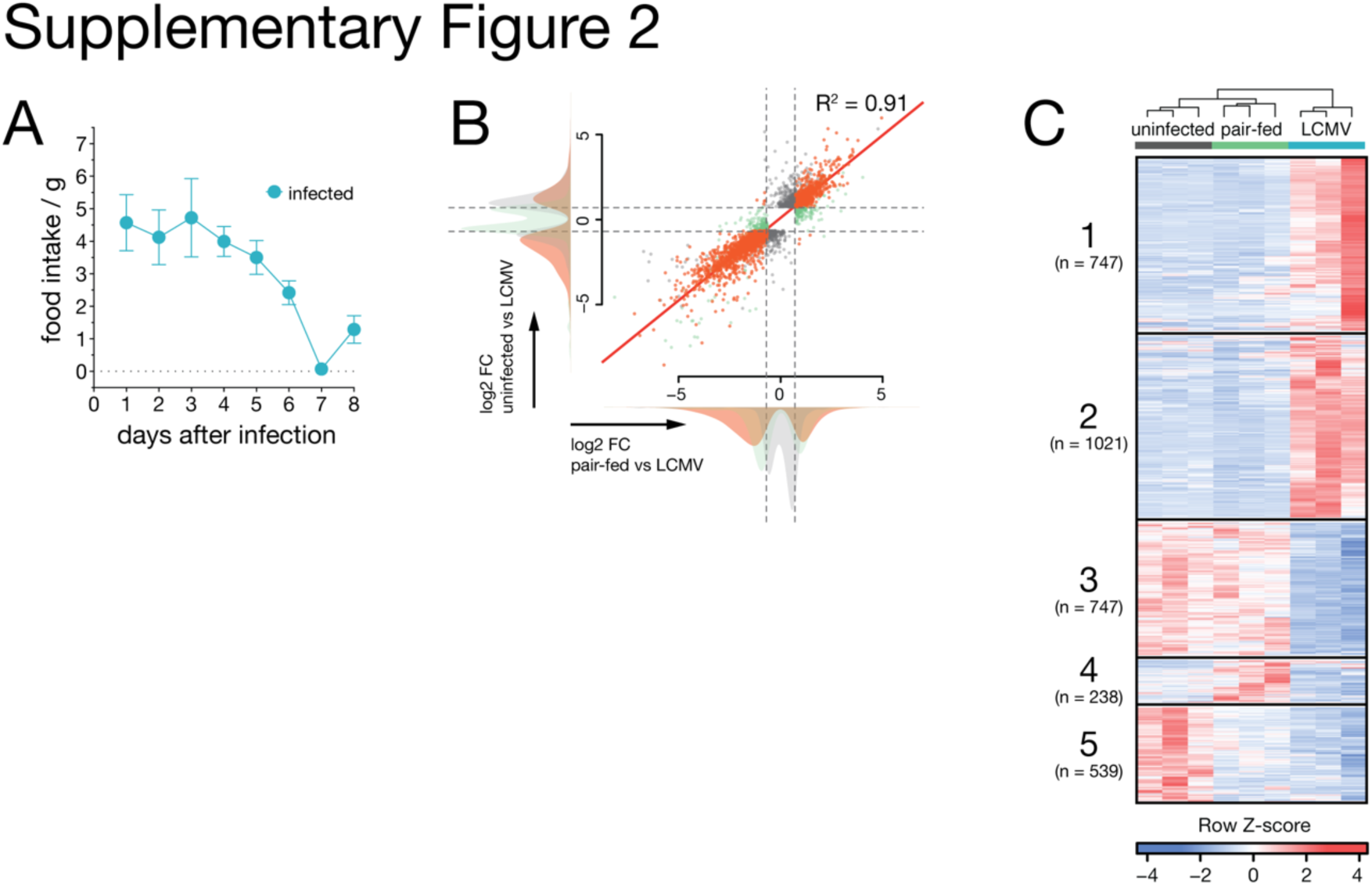
Transcriptome analyses of naïve and pair-fed compared to LCMV-infected animals. (A) Food intake of LCMV-infected animals. (B) Correlation of hepatic gene expression changes between naïve and pair-fed animals compared to LCMV infected animals. (C) Hierarchical clustering (FPKM, k-means, Pearson’s correlation) of significantly deregulated transcripts. Symbols represent the arithmetic mean ±S.E.M.

**Supplementary Figure S3:**
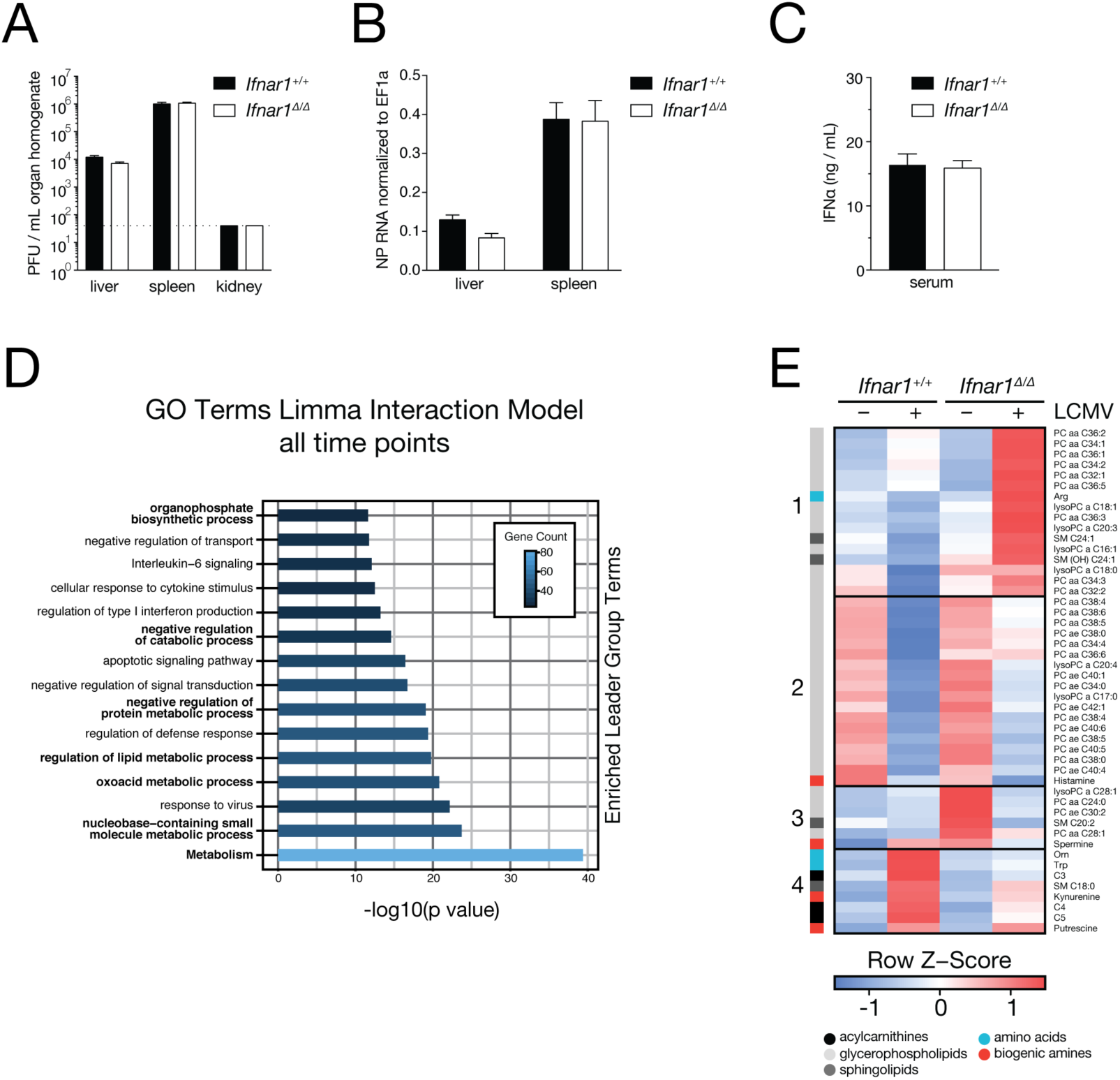
Transcriptome and systemic metabolome analyses of LCMV Cl13 infected Alb-Cre ERT2 Ifnar1fl/fl mice. (A) Viremia and (B) RNemia in organs, and (C) IFN-*α* serum levels of *Alb-Cre ERT2 Ifnar1^fl/fl^* (*Ifnar1^Δ/Δ^*) and *Ifnar1^+/+^* mice 1.5 days after infection. (D) Enriched GO terms on the union of differentially regulated genes (limma interaction model). (E) Significantly regulated serum metabolites in naïve and infected *Ifnar1^Δ/Δ^* and *Ifnar1^+/+^* animals (k-means, Pearson’s correlation). Symbols represent the arithmetic mean ±S.E.M. Dotted line implicates limit of detection. ns = not significant * P < 0.05 ** P < 0.01 *** P < 0.001(Student’s t-test).

**Supplementary Figure S4:**
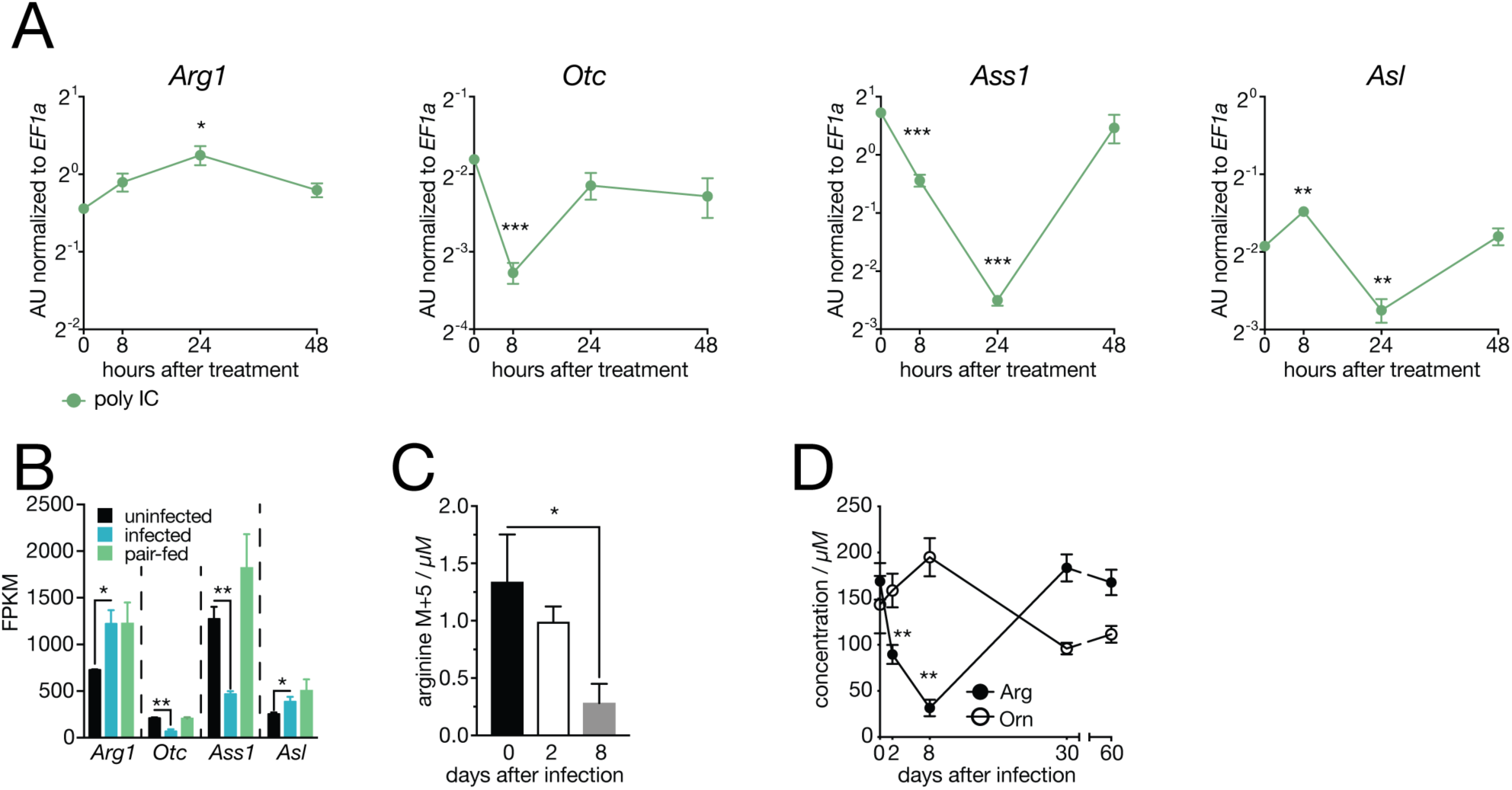
*Ifnar1* signaling regulates systemic arginine to ornithine ratio upon LCMV Cl13 infection. (A) Regulation of urea cycle-associated genes in liver tissue upon polyIC treatment of C57Bl/6J (wild type) mice (n = 3). (B) FPKM values of *Arg1*, *Otc*, *Ass1* and *Asl* of naïve, pair-fed and LCMV-infected (8dpi) wild type animals (n = 3). (C) Serum levels of ^13^C_5_ arginine upon LCMV clone 13 infection (n = 3-6). (D) Systemic arginine to ornithine concentrations upon LCMV infection (n = 8). Symbols represent the arithmetic mean ±S.E.M. ns = not significant * P < 0.05 ** P < 0.01 (Student’s t-test).

**Supplementary Figure S5:**
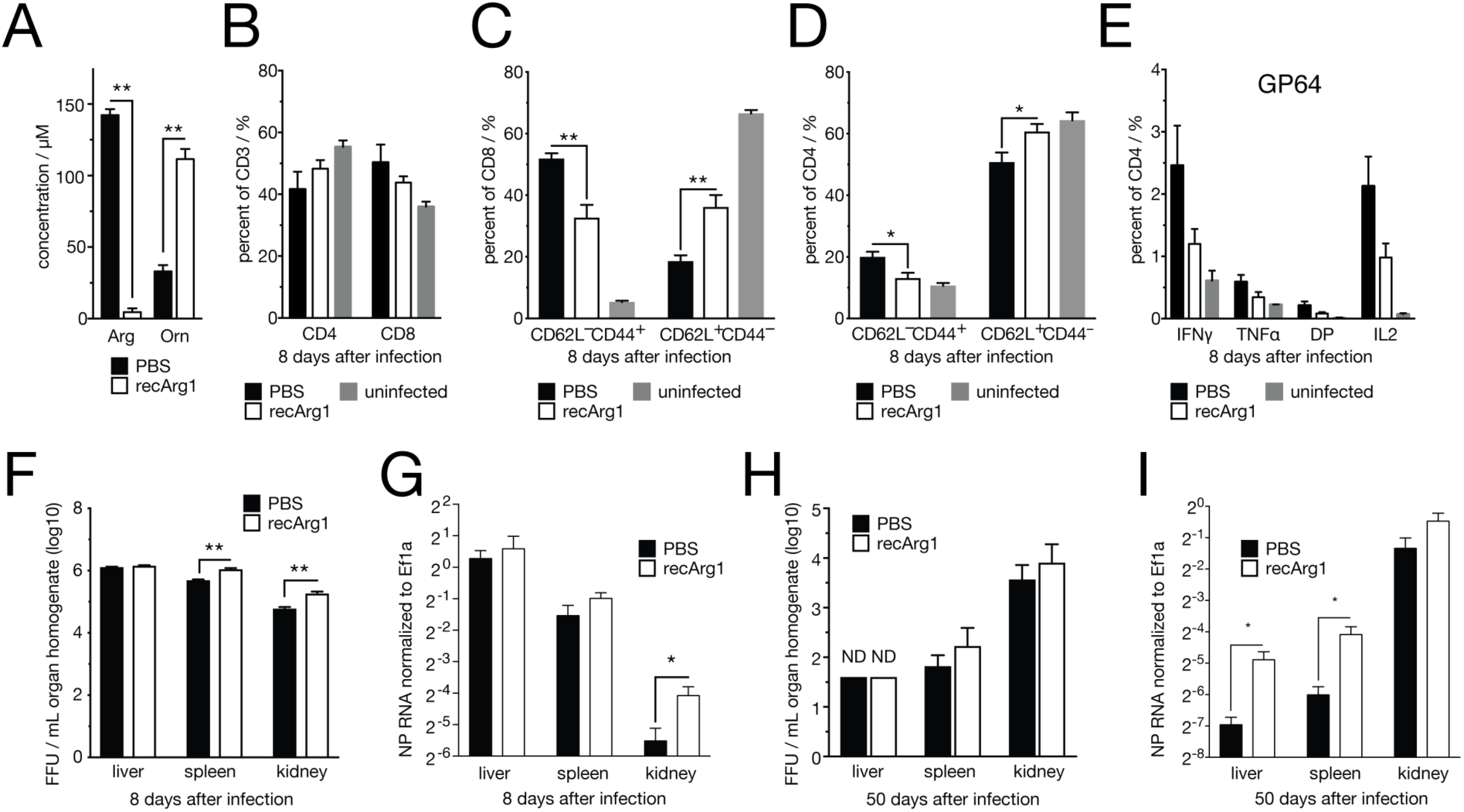
Arginase1 treatment does not affect splenic CD4 and CD8 T cell abundances or antiviral CD4 T cell responses and mildly affects organ viral load. (A) Serum arginine and ornithine levels in C57Bl/6J mice 2 days after treatment with pegylated recombinant human Arginase 1 (recArg1, BCT-100). (B) Percentage of CD4^+^ and CD8^+^ cells of CD3^+^ splenocytes at 8 days after LCMV clone 13 infection and recArg1 treatment. (C) CD62L^−^ CD44^+^ (effector) and CD62L^+^CD44^−^ (naïve) CD8 and (D) CD4 splenic T cells. (E) IFNγ, TNFα and IL2 production of LCMV GP64-specific splenic CD4 T cells 8 days after infection. (F) Viraemia and (G) RNemia of liver, spleen and kidney 8 days after infection. (H) Viraemia and (I) RNemia of liver, spleen and kidney 50 days after infection. Symbols represent the arithmetic mean ±S.E.M. n = 4-5. ns = not significant. ND = not detected. * P < 0.05 ** P < 0.01 (Student’s t-test).

**Supplementary Figure S6:**
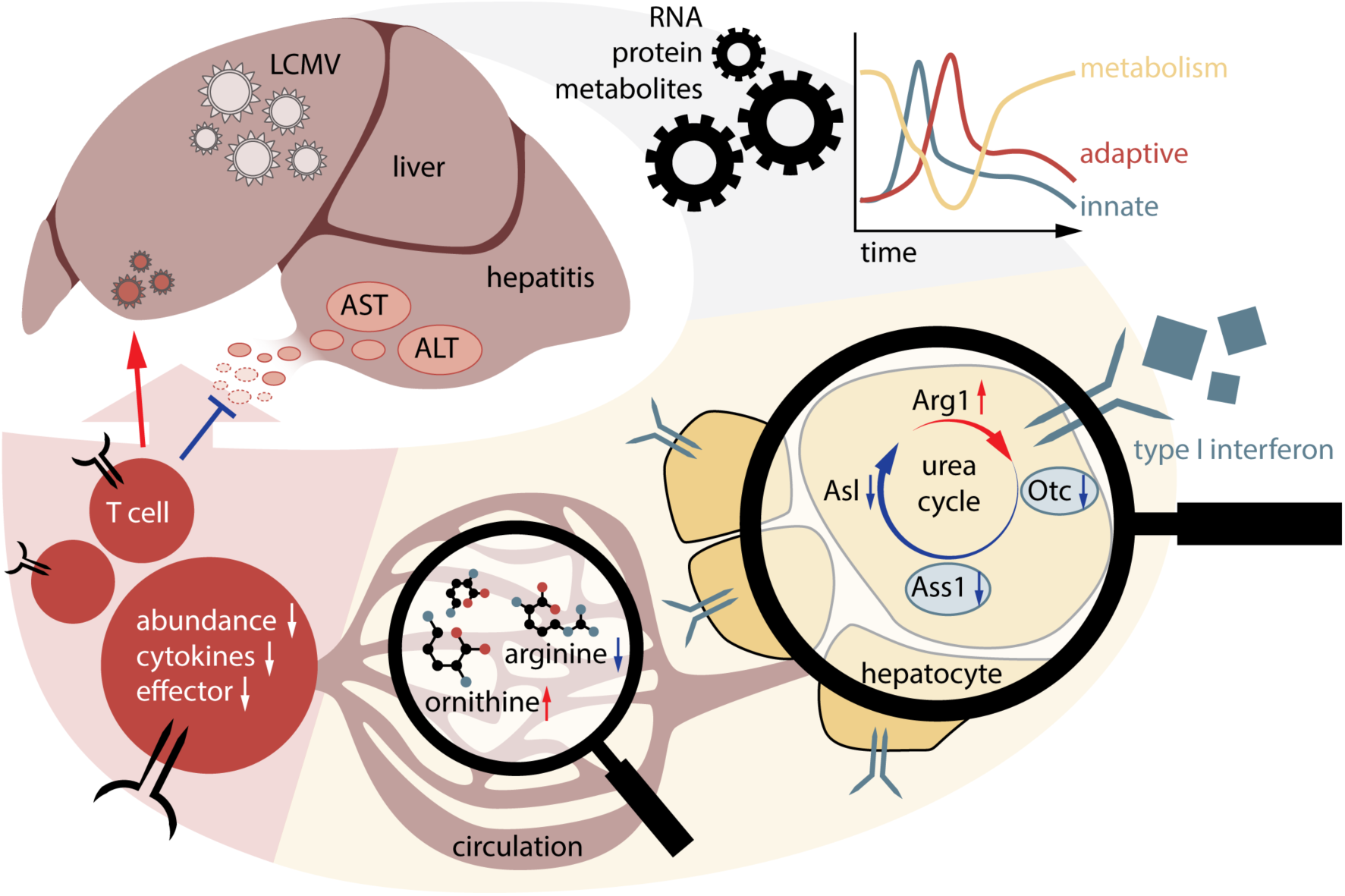
Working model. This graph depicts the working model of how chronic viral infection induces local changes of RNA, protein and metabolite levels in liver which, subsequently, translate into altered output of circulating metabolites. This metabolic reprogramming of the liver depends to a large extent on hepatocyte-intrinsic Ifnar1 signaling. Among the affected metabolic pathways, the urea cycle is affected by Ifnar1-dependent transcriptional regulation of its four enzymes *Arg1*, *Otc*, *Ass1* and *Asl*. Reduced levels of arginine and increased levels of ornithine in the serum suppress virus-specific CD8 T cell responses and reduce liver immunopathology.

**Table S1: Transcriptomic and proteomic data of LCMV-infected WT mice.**

**Table S2: Enrichment cluster analysis of differentially expressed genes.**

**Table S3: Metabolomics data of LCMV-infected mice.**

**Table S4: Transcriptomic and gene ontology enrichment data of pair-feeding experiment.**

**Table S5: Transcriptomic and gene ontology enrichment data of *Alb-Cre ERT2 Ifnar1* mice.**

## References

Adams, D.H., and Hubscher, S.G. (2006). Systemic viral infections and collateral damage in the liver. Am. J. Pathol. 168, 1057–1059.

Ah Mew, N., Simpson, K.L., Gropman, A.L., Lanpher, B.C., Chapman, K.A., and Summar, M.L. (1993). Urea Cycle Disorders Overview.

Baazim, H., Schweiger, M., Moschinger, M., Xu, H., Scherer, T., Popa, A., Gallage, S., Ali, S.A., Khamina, K., Kosack, L., et al. (2019). CD8 T cells induce cachexia in chronic viral infection. Nat. Immunol. In Press.

van den Berghe, G. (1991). The role of the liver in metabolic homeostasis: Implications for inborn errors of metabolism. J. Inherit. Metab. Dis. 14, 407–420.

Bhattacharya, A., Hegazy, A.N., Deigendesch, N., Kosack, L., Cupovic, J., Kandasamy, R.K., Hildebrandt, A., Merkler, D., Kühl, A.A., Vilagos, B., et al. (2015). Superoxide Dismutase 1 Protects Hepatocytes from Type I Interferon-Driven Oxidative Damage. Immunity 43, 974–986.

Bindea, G., Mlecnik, B., Hackl, H., Charoentong, P., Tosolini, M., Kirilovsky, A., Fridman, W.-H., Pagès, F., Trajanoski, Z., and Galon, J. (2009). ClueGO: a Cytoscape plug-in to decipher functionally grouped gene ontology and pathway annotation networks. Bioinformatics 25, 1091–1093.

Breitwieser, F.P., Müller, A., Dayon, L., Köcher, T., Hainard, A., Pichler, P., Schmidt-Erfurth, U., Superti-Furga, G., Sanchez, J.-C., Mechtler, K., et al. (2011). General statistical modeling of data from protein relative expression isobaric tags. J. Proteome Res. 10, 2758–2766.

Buck, M.D., Sowell, R.T., Kaech, S.M., and Pearce, E.L. (2017). Metabolic Instruction of Immunity. Cell 169, 570–586.

Casero, R.A., Murray Stewart, T., and Pegg, A.E. (2018). Polyamine metabolism and cancer: treatments, challenges and opportunities. Nat. Rev. Cancer 18, 681–695.

Chang, C.H., Qiu, J., O’Sullivan, D., Buck, M.D., Noguchi, T., Curtis, J.D., Chen, Q., Gindin, M., Gubin, M.M., Van Der Windt, G.J.W., et al. (2015). Metabolic Competition in the Tumor Microenvironment Is a Driver of Cancer Progression. Cell 162, 1229–1241.

Cheng, P.N.-M., Lam, T.-L., Lam, W.-M., Tsui, S.-M., Cheng, A.W.-M., Lo, W.-H., and Leung, Y.-C. (2007). Pegylated recombinant human arginase (rhArg-peg5,000mw) inhibits the in vitro and in vivo proliferation of human hepatocellular carcinoma through arginine depletion. Cancer Res. 67, 309–317.

Crispe, I.N. (2016). Hepatocytes as Immunological Agents. J. Immunol. 196, 17–21.

Dumas, M.-E., Kinross, J., and Nicholson, J.K. (2014). Metabolic phenotyping and systems biology approaches to understanding metabolic syndrome and fatty liver disease. Gastroenterology 146, 46–62.

Feun, L.G., Marini, A., Walker, G., Elgart, G., Moffat, F., Rodgers, S.E., Wu, C.J., You, M., Wangpaichitr, M., Kuo, M.T., et al. (2012). Negative argininosuccinate synthetase expression in melanoma tumours may predict clinical benefit from arginine-depleting therapy with pegylated arginine deiminase. Br. J. Cancer 106, 1481–1485.

Geiger, R., Rieckmann, J.C., Wolf, T., Basso, C., Feng, Y., Fuhrer, T., Kogadeeva, M., Picotti, P., Meissner, F., Mann, M., et al. (2016). L-Arginine Modulates T Cell Metabolism and Enhances Survival and Anti-tumor Activity. Cell 167, 829–842.e13.

Guidotti, L.G., Borrow, P., Brown, a, McClary, H., Koch, R., and Chisari, F.V.C.-P. (1999). Noncytopathic clearance of lymphocytic choriomeningitis virus from the hepatocyte. J Exp Med 189, 1555–64 ST-Noncytopathic clearance of lymphocyt.

Hotamisligil, G.S. (2017). Inflammation, metaflammation and immunometabolic disorders. Nature 542, 177–185.

Jenne, C.N., and Kubes, P. (2013). Immune surveillance by the liver. Nat. Immunol. 14, 996–1006.

Jha, A.K., Huang, S.C.C., Sergushichev, A., Lampropoulou, V., Ivanova, Y., Loginicheva, E., Chmielewski, K., Stewart, K.M., Ashall, J., Everts, B., et al. (2015). Network integration of parallel metabolic and transcriptional data reveals metabolic modules that regulate macrophage polarization. Immunity 42, 419–430.

Johnson, M.O., Wolf, M.M., Madden, M.Z., Andrejeva, G., Sugiura, A., Contreras, D.C., Maseda, D., Liberti, M. V., Paz, K., Kishton, R.J., et al. (2018). Distinct Regulation of Th17 and Th1 Cell Differentiation by Glutaminase-Dependent Metabolism. Cell 1–16.

Kanehisa, M., and Goto, S. (2000). KEGG: kyoto encyclopedia of genes and genomes. Nucleic Acids Res. 28, 27–30.

Kotas, M.E., and Medzhitov, R. (2015). Homeostasis, inflammation, and disease susceptibility. Cell 160, 816–827.

Krzywinski, M., Schein, J., Birol, I., Connors, J., Gascoyne, R., Horsman, D., Jones, S.J., and Marra, M.A. (2009). Circos: an information aesthetic for comparative genomics. Genome Res. 19, 1639–1645.

Lalazar, G., and Ilan, Y. (2014). Viral Diseases of the Liver. In Liver Immunology, (Cham: Springer International Publishing), pp. 159–171.

Lam, T.L., Wong, G.K.Y., Chow, H.Y., Chong, H.C., Chow, T.L., Kwok, S.Y., Cheng, P.N.M., Wheatley, D.N., Lo, W.H., and Leung, Y.C. (2011). Recombinant human arginase inhibits the in vitro and in vivo proliferation of human melanoma by inducing cell cycle arrest and apoptosis. Pigment Cell Melanoma Res. 24, 366–376.

Law, C.W., Chen, Y., Shi, W., and Smyth, G.K. (2014). voom: Precision weights unlock linear model analysis tools for RNA-seq read counts. Genome Biol. 15, R29.

Lee, J.S., Adler, L., Karathia, H., Carmel, N., Rabinovich, S., Auslander, N., Keshet, R., Stettner, N., Silberman, A., Agemy, L., et al. (2018). Urea Cycle Dysregulation Generates Clinically Relevant Genomic and Biochemical Signatures. Cell 1–12.

Li, H., Wang, L., Yan, X., Liu, Q., Yu, C., Wei, H., Li, Y., Zhang, X., He, F., and Jiang, Y. (2011). A proton nuclear magnetic resonance metabonomics approach for biomarker discovery in nonalcoholic fatty liver disease. J. Proteome Res. 10, 2797–2806.

Li, L., Mao, Y., Zhao, L., Li, L., Wu, J., Zhao, M., Du, W., Yu, L., and Jiang, P. (2019). p53 regulation of ammonia metabolism through urea cycle controls polyamine biosynthesis. Nature.

Liao, Y., Smyth, G.K., and Shi, W. (2014). featureCounts: an efficient general purpose program for assigning sequence reads to genomic features. Bioinformatics 30, 923–930.

Ma, E.H., Bantug, G., Griss, T., Condotta, S., Johnson, R.M., Samborska, B., Mainolfi, N., Suri, V., Guak, H., Balmer, M.L., et al. (2017). Serine Is an Essential Metabolite for Effector T Cell Expansion. Cell Metab. 25, 345–357.

McNab, F., Mayer-Barber, K., Sher, A., Wack, A., and O’Garra, A. (2015). Type I interferons in infectious disease. Nat. Rev. Immunol. 15, 87–103.

Medzhitov, R. (2008). Origin and physiological roles of inflammation. Nature 454, 428–435.

Meijer, A.J., Lamers, W.H., and Chamuleau, R.A. (1990). Nitrogen metabolism and ornithine cycle function. Physiol. Rev. 70, 701–748.

Miyajima, M., Zhang, B., Sugiura, Y., Sonomura, K., Guerrini, M.M., Tsutsui, Y., Maruya, M., Vogelzang, A., Chamoto, K., Honda, K., et al. (2017). Metabolic shift induced by systemic activation of T cells in PD-1-deficient mice perturbs brain monoamines and emotional behavior. Nat. Immunol. 18, 1342–1352.

Mostafavi, S., Yoshida, H., Moodley, D., Leboité, H., Rothamel, K., Raj, T., Ye, C.J., Chevrier, N., Zhang, S.Y., Feng, T., et al. (2016). Parsing the Interferon Transcriptional Network and Its Disease Associations. Cell 164, 564–578.

Mounce, B.C., Cesaro, T., Moratorio, G., Hooikaas, P.J., Yakovleva, A., Werneke, S.W., Smith, E.C., Poirier, E.Z., Simon-Loriere, E., Prot, M., et al. (2016). Inhibition of Polyamine Biosynthesis Is a Broad-Spectrum Strategy against RNA Viruses. J. Virol. 90, 9683–9692.

Murray, P.J. (2015). Amino acid auxotrophy as a system of immunological control nodes. Nat. Immunol. 17, 132–139.

Norata, G.D., Caligiuri, G., Chavakis, T., Matarese, G., Netea, M.G., Nicoletti, A., O’Neill, L.A.J., and Marelli-Berg, F.M. (2015). The Cellular and Molecular Basis of Translational Immunometabolism. Immunity 43, 421–434.

O’Neill, L.A.J., and Pearce, E.J. (2016). Immunometabolism governs dendritic cell and macrophage function. J. Exp. Med. 213, 15–23.

Okin, D., and Medzhitov, R. (2012). Evolution of inflammatory diseases. Curr. Biol. 22, R733–40.

Pallett, L.J., Gill, U.S., Quaglia, A., Sinclair, L. V, Jover-Cobos, M., Schurich, A., Singh, K.P., Thomas, N., Das, A., Chen, A., et al. (2015). Metabolic regulation of hepatitis B immunopathology by myeloid-derived suppressor cells. Nat. Med. 21, 591–600.

Pantel, A., Teixeira, A., Haddad, E., Wood, E.G., Steinman, R.M., and Longhi, M.P. (2014). Direct Type I IFN but Not MDA5/TLR3 Activation of Dendritic Cells Is Required for Maturation and Metabolic Shift to Glycolysis after Poly IC Stimulation. PLoS Biol. 12, e1001759.

Pearce, E.L., and Pearce, E.J. (2013). Metabolic pathways in immune cell activation and quiescence. Immunity 38, 633–643.

Pearce, E.L., Walsh, M.C., Cejas, P.J., Harms, G.M., Shen, H., Wang, L.-S., Jones, R.G., and Choi, Y. (2009). Enhancing CD8 T-cell memory by modulating fatty acid metabolism. Nature 460, 103–107.

Poillet-Perez, L., Xie, X., Zhan, L., Yang, Y., Sharp, D.W., Hu, Z.S., Su, X., Maganti, A., Jiang, C., Lu, W., et al. (2018). Autophagy maintains tumour growth through circulating arginine. Nature 563, 569–573.

Protzer, U., Maini, M.K., and Knolle, P.A. (2012). Living in the liver: hepatic infections. Nat. Rev. Immunol. 12, 201–213.

Rabinovich, S., Adler, L., Yizhak, K., Sarver, A., Silberman, A., Agron, S., Stettner, N., Sun, Q., Brandis, A., Helbling, D., et al. (2015). Diversion of aspartate in ASS1-deficient tumours fosters de novo pyrimidine synthesis. Nature 527, 379–383.

Racanelli, V., and Rehermann, B. (2006). The liver as an immunological organ. Hepatology 43, S54–62.

Rehermann, B., and Nascimbeni, M. (2005). Immunology of hepatitis B virus and hepatitis C virus infection. Nat. Rev. Immunol. 5, 215–229.

Sanchez, K.K., Chen, G.Y., Schieber, A.M.P., Redford, S.E., Shokhirev, M.N., Leblanc, M., Lee, Y.M., and Ayres, J.S. (2018). Cooperative Metabolic Adaptations in the Host Can Favor Asymptomatic Infection and Select for Attenuated Virulence in an Enteric Pathogen. Cell 175, 146–158.e15.

De Santo, C., Cheng, P., Beggs, A., Egan, S., Bessudo, A., and Mussai, F. (2018). Metabolic therapy with PEG-arginase induces a sustained complete remission in immunotherapy-resistant melanoma. J. Hematol. Oncol. 11, 1–5.

Schoggins, J.W., Wilson, S.J., Panis, M., Murphy, M.Y., Jones, C.T., Bieniasz, P., and Rice, C.M. (2011). A diverse range of gene products are effectors of the type i interferon antiviral response. Nature 472, 481–485.

Sinclair, L. V., Rolf, J., Emslie, E., Shi, Y.B., Taylor, P.M., and Cantrell, D.A. (2013). Control of amino-acid transport by antigen receptors coordinates the metabolic reprogramming essential for T cell differentiation. Nat. Immunol. 14, 500–508.

Smyth, G.K. (2004). Linear models and empirical bayes methods for assessing differential expression in microarray experiments. Stat. Appl. Genet. Mol. Biol. 3, Article3.

Soga, T., Sugimoto, M., Honma, M., Mori, M., Igarashi, K., Kashikura, K., Ikeda, S., Hirayama, A., Yamamoto, T., Yoshida, H., et al. (2011). Serum metabolomics reveals γ-glutamyl dipeptides as biomarkers for discrimination among different forms of liver disease. J. Hepatol. 55, 896–905.

St John-Williams, L., Blach, C., Toledo, J.B., Rotroff, D.M., Kim, S., Klavins, K., Baillie, R., Han, X., Mahmoudiandehkordi, S., Jack, J., et al. (2017). Targeted metabolomics and medication classification data from participants in the ADNI1 cohort. Sci. Data 4, 170140.

Uhlen, M., Fagerberg, L., Hallstrom, B.M., Lindskog, C., Oksvold, P., Mardinoglu, A., Sivertsson, A., Kampf, C., Sjostedt, E., Asplund, A., et al. (2015). Tissue-based map of the human proteome. Science (80-.). 347, 1260419–1260419.

Van De Velde, L.A., and Murray, P.J. (2016). Proliferating helper T cells require rictor/mTORC2 complex to integrate signals from limiting environmental amino acids. J. Biol. Chem. 291, 25815–25822.

Virgin, H.W., Wherry, E.J., and Ahmed, R. (2009). Redefining Chronic Viral Infection. Cell 138, 30–50.

Watford, M. (2003). The urea cycle: Teaching intermediary metabolism in a physiological setting. Biochem. Mol. Biol. Educ. 31, 289–297.

Wu, D., Sanin, D.E., Everts, B., Chen, Q., Qiu, J., Buck, M.D., Patterson, A., Smith, A.M., Chang, C.-H., Liu, Z., et al. (2016). Type 1 Interferons Induce Changes in Core Metabolism that Are Critical for Immune Function. Immunity 44, 1325–1336.

York, A.G., Williams, K.J., Argus, J.P., Zhou, Q.D., Brar, G., Vergnes, L., Gray, E.E., Zhen, A., Wu, N.C., Yamada, D.H., et al. (2015). Limiting Cholesterol Biosynthetic Flux Spontaneously Engages Type I IFN Signaling. Cell 163, 1716–1729.

Zehn, D., and Wherry, E.J. (2015). Immune Memory and Exhaustion: Clinically Relevant Lessons from the LCMV Model. Adv. Exp. Med. Biol. 850, 137–152.

Zhou, Z., Xu, M.-J., and Gao, B. (2015). Hepatocytes: a key cell type for innate immunity. Cell. Mol. Immunol. 1–15.

Zinkernagel, R.M., Haenseler, E., Leist, T., Cerny, A., Hengartner, H., and Althage, A. (1986). T cell-mediated hepatitis in mice infected with lymphocytic choriomeningitis virus. Liver cell destruction by H-2 class I-restricted virus-specific cytotoxic T cells as a physiological correlate of the 51Cr-release assay? J Exp Med 164, 1075–1092.

